# A cytological and functional framework of the meiotic spindle assembly checkpoint in *Arabidopsis thaliana*

**DOI:** 10.1101/2023.05.26.542430

**Authors:** Konstantinos Lampou, Franziska Böwer, Shinichiro Komaki, Maren Köhler, Arp Schnittger

**Affiliations:** Department of Developmental Biology, Institute for Plant Sciences and Microbiology, University of Hamburg, 22609 Hamburg, Germany; Graduate School of Science and Technology, Nara Institute of Science and Technology, Nara, 630-0192 Japan

**Keywords:** cell division, live imaging, meiosis, microtubule, spindle assembly checkpoint

## Abstract

The spindle assembly checkpoint (SAC) is a surveillance mechanism active during metaphase to prevent aneuploidy. The SAC is especially important during meiosis to maintain genome stability over generations and sustain fertility. However, despite its crucial role for reproduction and breeding, little is known about the plant meiotic SAC. Here, we present a cytological and functional framework of the SAC in male meiocytes of *Arabidopsis thaliana*. Using live-cell imaging, we have dissected the temporal association of SAC components with the kinetochore and have identified the three conserved kinases BMF1, MPS1 and AURORA as crucial regulators of the loading of BMF3 to kinetochores. Functionally characterizing core SAC components, we found that BUB3.3 has a predominant and previously not recognized role in chromosome congression. We suggest that BUB3.3 is involved in efficient kinetochore-microtubule interactions. Furthermore, the meiotic SAC is only active for a limited time under severe microtubule destabilizing conditions leading to the hypothesis that the relaxed nature of the meiotic SAC is a gateway to polyploidization and hence might contribute to genome evolution in plants.

## Introduction

Sexual reproduction requires the generation of gametes, which have only half of the chromosome number in comparison to their parental organism, so that after fertilization the full chromosome set is restored. The reduction of chromosome number relies on elaborated segregation events accomplished in meiosis.

The spindle assembly checkpoint (SAC) is a safe-guarding mechanism blocking the metaphase-to-anaphase transition in meiosis (as well as in mitosis) until all kinetochores are correctly attached to spindle microtubules emanating from opposing poles. Thus, the SAC ensures error-free chromosome segregation and consequently, defects in the SAC cause severe developmental maladies such as cancer and infertility. For this reason, the SAC has been well-explored in mammals. There, Mps1 is the conductor of SAC assembly (London & Biggins, 2014). Mps1 is recruited to unattached kinetochores where it phosphorylates several kinetochore components (Hiruma et al., 2015; Ji et al., 2015). These are recognized by Bub3 which further recruits Bub1 and BubR1 (Primorac et al., 2013; Vleugel et al., 2013, 2015). A Mad1-Mad2 heterotetramer binds then to Mps1-phosphorylated Bub1 (Ji et al., 2017; G. Zhang et al., 2017). Ultimately, the SAC produces a cytosolic inhibitor, the mitotic checkpoint complex (MCC), a tetrameric complex composed of BubR1, Bub3, Mad2 and Cdc20 (Sudakin et al., 2001). The MCC acts as a pseudosubstrate of the anaphase promoting complex/cyclosome (APC/C), a multi-subunit ubiquitin E3 ligase which licenses the metaphase-to-anaphase transition (Alfieri et al., 2016; Izawa & Pines, 2015; Lara-Gonzalez et al., 2011). Once all kinetochores are properly attached, the SAC is silenced.

Despite the overall similarities, there are kingdom-as well as species-specific features of the SAC. In our previous work, we studied the SAC in roots and functionally characterized the homologs of all SAC components in Arabidopsis revealing some conserved but also several unique features of the plant mitotic SAC (Komaki & Schnittger, 2017). A characteristic feature of SAC proteins is their transient recruitment to and association with the incorrectly attached kinetochore. Surprisingly, both, BMF1, one of the three Arabidopsis Bub1/Mad3 family members, and MPS1 localize on the kinetochore throughout the cell cycle. BMF1 was hypothesized to not be part of the SAC signaling cascade since the *bmf1* mutant is not hypersensitive to oryzalin, which prevents microtubule polymerization (Komaki & Schnittger, 2017). MPS1, on the other hand, plays a key role in the plant SAC by recruiting MAD2 to the kinetochore. BubR1 functions seem to be shared between the two other Bub1/Mad3 family members BMF2 and BMF3. BMF2 is probably part of the MCC as suggested by the presence of two KEN-boxes essential for BubR1 function (Lara-Gonzalez et al., 2011; Malureanu et al., 2009), while BMF3 moves onto the kinetochore during prometaphase and is the platform on which the MAD1-MAD2 complex assembles. Finally, there are three Bub3 homologs in Arabidopsis, of which only BUB3.3 appears to be involved in the SAC, while BUB3.1 and BUB3.2 localize to spindle microtubules and are involved in phragmoplast formation (Komaki & Schnittger, 2017; H. Zhang et al., 2018). Reminiscent of yeast, the plant SAC is redundant for survival under non-stress conditions (Hoyt et al., 1991; Komaki & Schnittger, 2017; Li & Murray, 1991). Under severe microtubule destabilizing conditions, plant cells shut down the SAC after about 2 hours and rebuild a nuclear envelope, thus generating a cell with a duplicated chromosome set.

Previous studies have indicated that there are differences between the meiotic and mitotic SAC (Gorbsky, 2015). Despite of its importance for the viability of the progeny, little is currently known about the plant meiotic SAC. At the same time, meiocytes in Arabidopsis could be a powerful model system to study the SAC since meiotic divisions are highly synchronous in anthers, the male sexual organs (Prusicki et al., 2019). Moreover, due to the mono-orientation of kinetochores of the sister chromatids in meiosis I, it is presumably easier to detect and follow SAC components in comparison to the single sister chromatid kinetochores in mitosis.

Here, we have set out to generate a cell-biological and functional framework of the SAC in meiosis of Arabidopsis. Using live-cell imaging of male meiocytes, we first explore dynamics of the SAC assembly and disassembly on kinetochore and explore which kinases likely play part in this. Interestingly, while our work shows that BMF1 does play a role in the plant SAC, we also reveal that BUB3.3 does not seem to be a *bona fide* SAC component but is rather involved in the formation of stable, bipolar kinetochore-microtubule attachments.

## Results and Discussion

### Spatiotemporal localization and step-wise (dis-)assembly of SAC components in meiosis I

Using the functional genomic reporters developed previously (Komaki & Schnittger, 2017), we explored the localization of SAC components in space and time in meiosis I (Fig. 1) and meiosis II (Fig. EV1). TUA5 fused to TagRFP was used to identify meiotic stages based on microtubule organization patterns (Prusicki et al., 2019). Importantly, the TUA5 reporter also allowed us to visualize the spindle and monitor the timing of <u>n</u>uclear <u>e</u>nvelope <u>b</u>reakdown (NEB) until <u>a</u>naphase <u>o</u>nset (AO).

**Figure 1.**
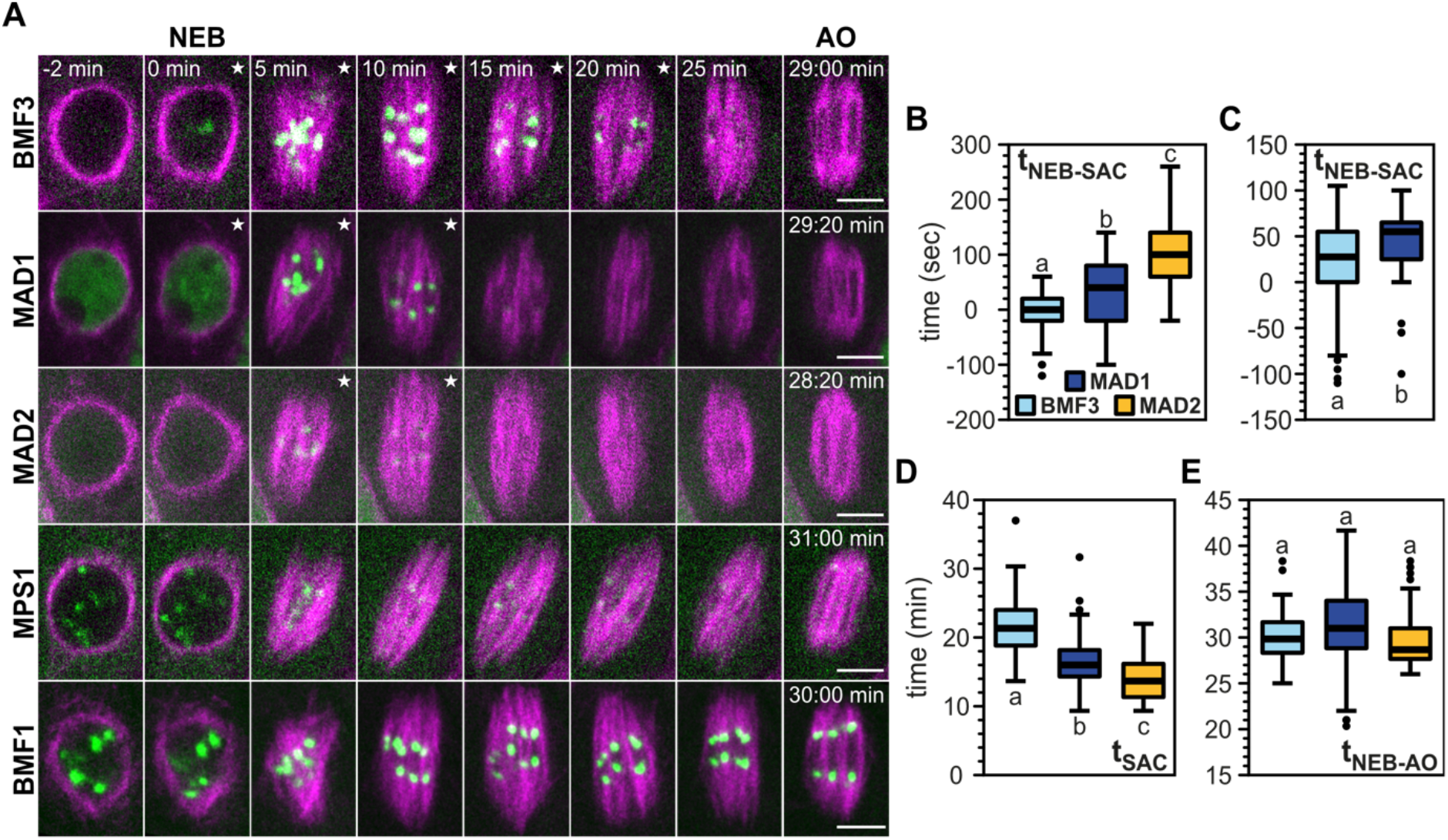
Spatiotemporal characterization of functional reporters of core SAC components in meiosis I. **(A)** Localization of core SAC proteins BMF3, MAD1, MAD2, MPS1 and BMF1 fused to GFP (green) together with TUA5 fused to TagRFP (magenta) decorating microtubules in metaphase I. MPS1 and BMF1 are localized at the kinetochore throughout metaphase and hence, they were excluded from further quantification in **(B)**-**(E)**. **(B,C)** Quantification of time needed for the kinetochore signal to appear relative to NEB. **(D)** Quantification of duration of the kinetochore signal for each reporter. To exemplify how we timed the kinetochore localization, the relevant frames are decorated with a white star (★) on the upper right corner. **(E)** Quantification of metaphase I duration defined as the time between NEB and AO for each reporter line. Movies were recorded with a frame rate of 20 **(A,B,D and E)** or 5 **(C)** seconds. For **(B)**, **(D)** and **(E)**, we used the same set of movies and quantified 44-84 cells from 8-9 different anthers for each reporter line. The values shown in **(C)** are from a different set of movies and we quantified 70-76 cells from 15-16 anthers for each reporter line. Different letters indicate significance (P < 0.01; pairwise comparison using Kruskal-Wallis test for independent samples with Bonferroni correction for multiple tests). Scale bars: 5 µm.

With the exception of the cytosolic localization of BMF2 and BUB3.3 (Fig. EV1A and Video 1), all SAC reporters exhibit a dotted appearance indicative of their localization at kinetochores (Fig. 1A and Video 2) similar to mitosis (Komaki & Schnittger, 2017). In contrast to human mitotic cells (Overlack et al., 2015; Stucke et al., 2002; Vleugel et al., 2015) but similar to mitosis in Arabidopsis, MPS1 and BMF1 localize on kinetochores already before NEB (Fig. 1A and Video 2).

As a next step towards a cytological framework of the SAC, we sought to quantify when the other SAC proteins appear on and disappear from the kinetochore relative to the NEB in meiosis I. With a frame rate of 20 seconds, we determined that BMF3 appears first on kinetochores after NEB, quickly followed by MAD1 and then MAD2, which became visible on average after ∼ 100 seconds after NEB (Fig. 1A, B). To better resolve the relative timing of BMF3 and MAD1, we then recorded movies with a frame rate of 5 seconds revealing that BMF3 needs on average ∼ 30 seconds and MAD1 ∼ 50 seconds to appear on the kinetochore (Fig. 1B and C). To assess the validity of our assays, we determined the duration of metaphase I in our reporter lines. Indeed, we did not observe any significant difference in the duration of NEB to AO (t_NEB-AO_) among all our reporter lines indicating that these reporters do not interfere with the progression of meiosis (Fig. 1E).

The disassembly of the kinetochore-bound SAC complex, determined by the disappearance of the dotty fluorescence signal, happened in the opposite order of the assembly, i.e., MAD2 disappeared first, i.e., ∼ 14 minutes after NEB, followed by MAD1 after ∼ 17 minutes and finally, BMF3 after ∼ 22 minutes. These observations suggest a strict hierarchy in SAC association with and disassociation from kinetochores (Fig. 1D).

Next, we checked the localization of the BMF3, MAD1 and MAD2 reporters in meiosis II (Fig. EV1B and Video 3). We did not observe any obvious difference between metaphase I and metaphase II. Hence, we will focus only on metaphase I from here onwards.

While the turnover dynamics of SAC proteins at the kinetochore have been studied extensively (Howell et al., 2004; Shah et al., 2004), to the best of our knowledge, however, there is no work in other model species showing the sequential association to and disassociation from the kinetochore of SAC components. Based on our observations, we conclude that the assembly and disassembly pattern reflect a general principle of SAC composition in Arabidopsis and likely other plant species.

### BMF1, MPS1 and AURORA kinases positively affect the timely recruitment of BMF3 to the kinetochore

The step-wise (dis-)assembly of the SAC argued for a strict regulation of protein interactions at the level of the kinetochore, possibly through post-translational modifications. Phosphorylation is well known to control the interaction of SAC components and finetune the strength of the SAC in mammals for instance by promoting the interaction of Bub3-Bub1 and Bub3-BubR1 dimers with Knl1 (preprint: Corno et al., 2022; Espert et al., 2014; Vleugel et al., 2013).

We have previously reported that the BMF3 localization is not affected in *bmf1* and *mps1* mutants on a qualitative level (Komaki & Schnittger, 2017). Given the results above, we wondered, however, whether this would hold true on a quantitative level. For this, we introgressed the BMF3 reporter into the *bmf1* and *mps1* backgrounds through crossing. When imaging these lines with identical settings, the BMF3 kinetochore signal in *bmf1* and *mps1* mutants appeared to be reduced (Fig. 2A, EV2A and Video 4). To quantify this effect, we measured the fluorescence intensity (FI) of BMF3 in the nucleus from NEB-AO. The BMF3 signal was normalized to the frame of the NEB and the maximum value for each cell was plotted (Fig. 2B). Our quantification uncovered a previously unnoticed effect of BMF1- and MPS1-dependent BMF3 recruitment to the kinetochore both regarding the protein amount (Fig. 2B) as well as the timing of recruitment (t_NEB-BMF3_) (Fig. 2C).

**Figure 2.**
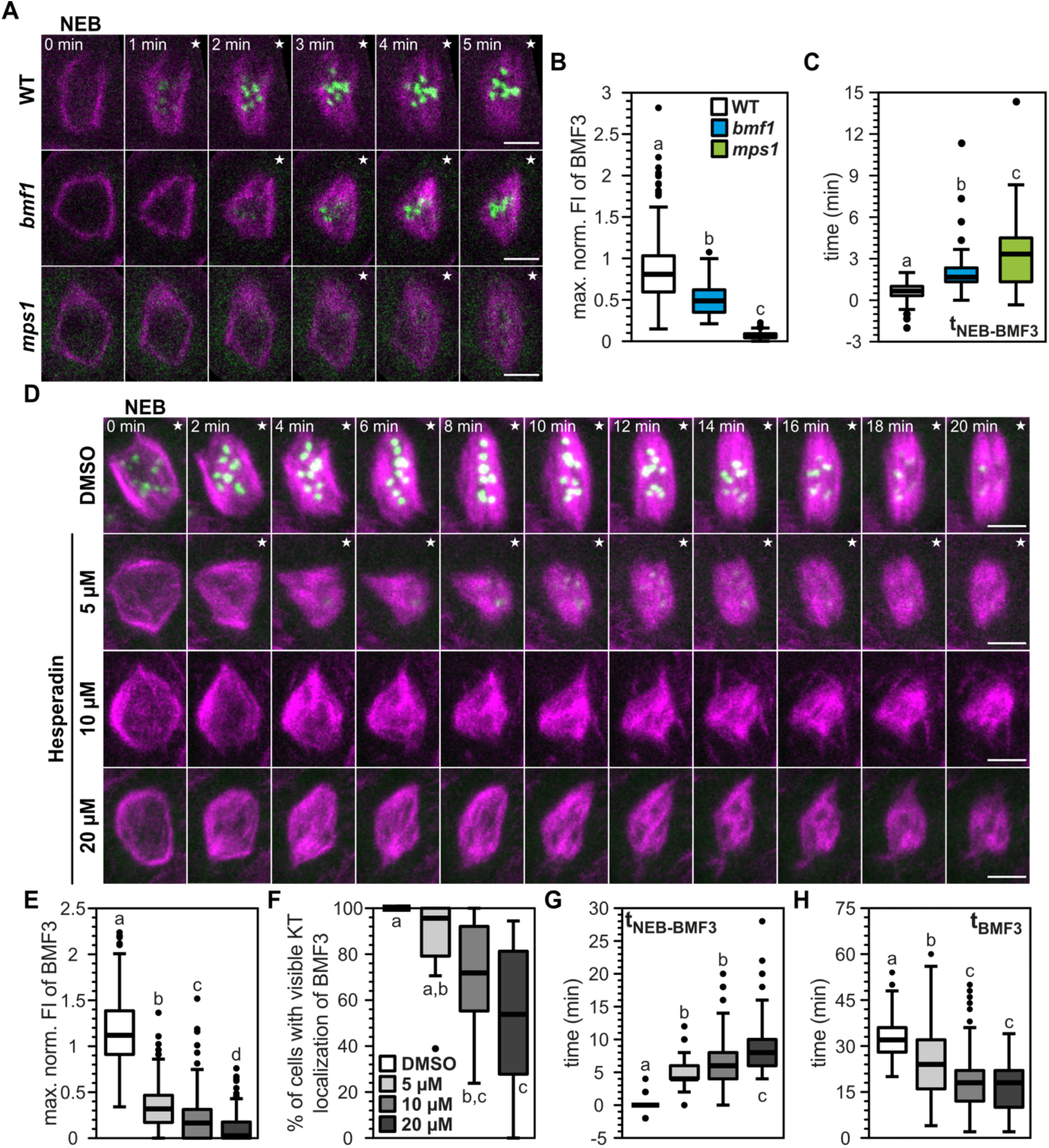
BMF1, MPS1 and AURORA kinases are involved in the kinetochore recruitment of BMF3 in metaphase I. **(A)** BMF3 recruitment in WT, *bmf1* and *mps1* in metaphase I. BMF3 is fused to GFP (green) and TUA5 to TagRFP (magenta). **(B)** Quantification of maximum fluorescence intensity (FI) of BMF3 during metaphase I normalized to the frame of NEB: (FI_max_-FI_NEB_)/FI_NEB_. **(C)** Quantification of time needed for the BMF3 kinetochore signal to appear relative to NEB. **(D)** Maximum projections of representative WT cells in metaphase I treated with 5, 10 or 20 µM hesperadin or the equivalent amount of DMSO. BMF3 is fused to GFP (green) and TUA5 to TagRFP (magenta). **(E)** Quantification of maximum FI of BMF3 during metaphase I normalized to the frame of NEB as in **(B)**. **(F)** Percentage of cells per anther with visible BMF3 on the kinetochore during metaphase I. **(G)** Quantification of time needed for the BMF3 kinetochore signal to appear relative to NEB. **(H)** Quantification of duration of the BMF3 kinetochore signal. The white star (★) on the upper right corner of the frames indicates BMF3 kinetochore localization. Movies were recorded with a frame rate of 20 seconds **(A-C)** or 2 minutes **(D-H)**. For **(A)-(C)**, we quantified 63-242 cells from 6-19 anthers per genotype. For **(E)-(H)**, we quantified 151-339 cells from 13-16 anthers per treatment. Different letters indicate significance (P < 0.01; pairwise comparison using Kruskal-Wallis test for independent samples with Bonferroni correction for multiple tests). Scale bars: 5 µm.

Additionally, we scored visually whether the BMF3 kinetochore signal appeared after NEB. While 100% of wild type (WT) and *bmf1* cells displayed a BMF3 kinetochore signal, the percentage of cells having visually recognizable kinetochore signal varied from 0% to a 100% in *mps1* anthers arguing for an especially prominent role of MPS1 in recruiting BMF3 to the kinetochore (Fig. EV2B). It is unclear to us, what might cause the large variation between *mps1* cells with and without a recognizable BMF3 kinetochore signal within a single anther as well as between anthers.

There is strong evidence in mammalian cells that Aurora B (AurB) activates the SAC not only through destabilization of incorrect kinetochore-microtubule attachments but also through directly phosphorylating central regulators of the SAC (Roy et al., 2022; Santaguida et al., 2011; Saurin et al., 2011). There are two families of AUR kinases in plants: α-AUR kinases with AUR1 and AUR2 and β-AUR kinases with AUR3 (Demidov et al., 2009). AUR3 is the closest homolog of the mammalian Aurora B (AurB). AUR1 and AUR2 localize to the spindle while AUR3 localizes to the centromere during metaphase (Demidov et al., 2009; Komaki et al., 2020; Komaki & Schnittger, 2017). Hence, we hypothesized that the AUR family might influence the plant SAC assembly and/or function in parallel to MPS1 and BMF1.

To test this hypothesis, we decided to use hesperadin, an AurB-specific inhibitor in mammals (Hauf et al., 2003), that has been shown to inhibit at least two of the three plant AURs, albeit AUR1 is inhibited at a much lower concentration than AUR3 (Demidov et al., 2009; Kurihara et al., 2006). We treated flowers with 5, 10 or 20 µM hesperadin or the equivalent amount of DMSO (Fig. 2D, EV2D and Video 5). Similar to *bmf1* and *mps1*, increasing concentration of hesperadin resulted in ever decreasing maximum FI of BMF3 kinetochore signal (Fig. 2E) and fewer cells having a visible BMF3 kinetochore signal (Fig. 2F). Furthermore, the BMF3 kinetochore signal appeared at an increasingly later time after NEB (Fig. 2G) and persisted for shorter (t_BMF3_) with higher concentrations of hesperadin (Fig. 2H). In addition, the spindle architecture became severely affected with an increasing concentration of hesperadin (Fig. 2D) while the metaphase I duration – albeit shorter than in the DMSO control – remained unchanged with an increasing concentration of hesperadin (Fig. EV2C).

In conclusion, the major kinases BMF1, MPS1 and the AURORA family are orchestrating the timely and sufficient assembly of the SAC by regulating either directly or indirectly BMF3 localization. Given the AUR1/2/3 localization patterns, most probably AUR3 is responsible for recruiting BMF3 to the kinetochore. Considering that AurB and Mps1 both regulate each other (Jelluma et al., 2008; Santaguida et al., 2011; Saurin et al., 2011; Van Der Waal et al., 2012) and that AurB also regulates Bub1 in mammalian cells (Roy et al., 2022), it would be interesting to untangle in the future whether AUR3 is regulating MPS1 and BMF1 in plants or vice versa.

### Putting the meiotic SAC to a test

As a next step, we decided to test whether the meiotic SAC is functional, considering that SAC mutants in Arabidopsis do not exhibit any severe developmental defects (Komaki & Schnittger, 2017). First, we compared the duration of the BMF3 kinetochore signal, which from here onwards we use as a proxy for an active SAC, and the metaphase I duration in the WT versus *mad2* (Fig. 3A, B and Video 6). In support of an active meiotic SAC, the BMF3 kinetochore signal persisted for shorter in *mad2* (Fig. 3B) and correspondingly, the duration of metaphase I was ∼ 30% shorter than in the WT.

**Figure 3.**
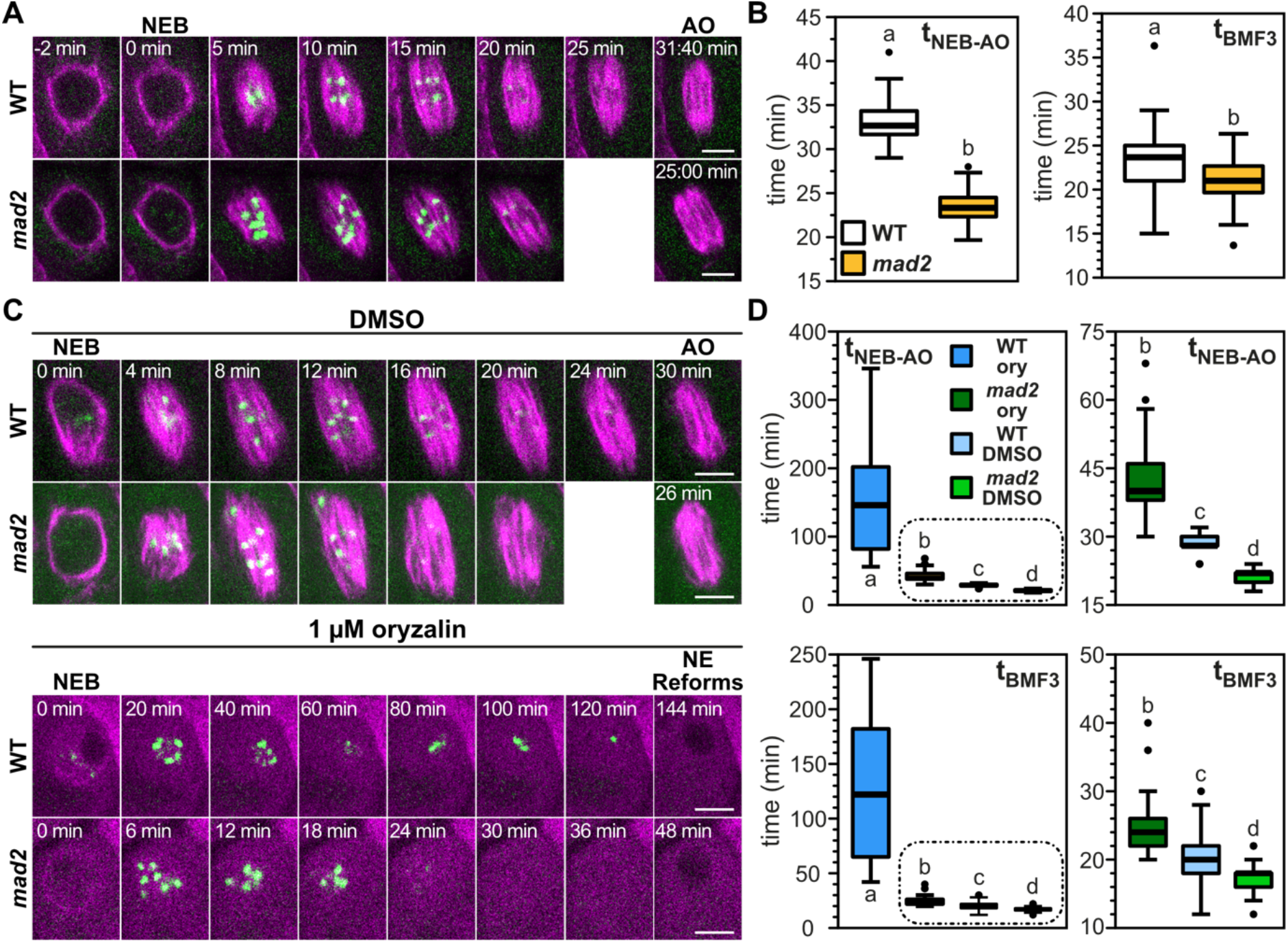
The meiotic SAC is functional but permissive. **(A)** BMF3 kinetochore signal and metaphase duration in representative WT and *mad2* cells in metaphase I. BMF3 is fused to GFP (green) and TUA5 to TagRFP (magenta). **(B)** Quantification of metaphase I (left) and of BMF3 kinetochore signal (right) duration in the WT and *mad2*. **(C)** BMF3 kinetochore signal and metaphase duration in representative WT and *mad2* cells in metaphase I when treated with 1 µM oryzalin or the equivalent amount of DMSO. Note that in the oryzalin treatment the SAC gets silenced and the NE reforms. **(D)** Quantification of metaphase I (up) and of BMF3 kinetochore signal (down) duration in the WT and *mad2*. The encircled box plots in the graphs on the left side, are shown larger on the right side. 2D movies were recorded with a frame rate of 20 seconds **(A,B)** or 2 minutes **(C,D)**. For **(B)**, we quantified 89-122 cells from 16-20 anthers per genotype. For **(D)**, we quantified 63-133 cells from 13-22 anthers per treatment and genotype. Different letters indicate significance (P < 0.01; pairwise comparison using Kruskal-Wallis test for independent samples with Bonferroni correction for multiple tests). Scale bars: 5 µm.

A prolonged SAC-dependent metaphase arrest can be caused in most animal and fungal cells by drugs that depress microtubule polymerization, such as oryzalin (Brito & Rieder, 2006; Rieder & Maiato, 2004). These drugs either weaken or abolish the formation of stable and correct microtubule-kinetochore attachments, thus keeping the SAC in an active state. Supporting an active role of the SAC in meiosis, treatment with oryzalin led to a substantial increase in metaphase I duration in the WT but not in *mad2* mutants (Fig 3C, D and Video 7).

Many animal and fungal cells that arrest in metaphase might either become apoptotic or exit mitosis without satisfying the SAC, a process termed “mitotic slippage” or “checkpoint adaptation” often causing aneuploidy (Rieder & Maiato, 2004). In contrast, slippage could not be detected in previous work analyzing Arabidopsis root cells (Komaki & Schnittger, 2017). Considering that there are noticeable differences between the mitotic and meiotic SAC activity in other species (Gorbsky, 2015), we monitored the SAC duration of oryzalin treated flowers and found that the metaphase I arrest in the WT was on average ∼ 2 hours long and could never be sustained for longer than 6 hours (Fig. 3D). Correspondingly, the SAC was shut off within ∼ 2 hours on average as indicated by the disappearance of the BMF3 kinetochore signal and eventually a NE reformed. This is reminiscent of the situation in mitosis of Arabidopsis where an active SAC can be maintained for up to approximately 90 minutes (Komaki & Schnittger, 2017). Thus, SAC silencing followed by NE reformation appear to be general principles of the Arabidopsis response to high concentration of MT destabilizing drugs and possibly that of other plants. Surprisingly, all male meiocytes underwent a second NEB event marking the onset of meiosis II, indicating that NEB dynamics are independent of microtubules (Fig. EV3A and Video 8). In addition, we observed that interphase duration decreased from ∼ 53-55 minutes in DMSO treated cells to ∼ 38-39 minutes in cells treated with oryzalin. Considering that this was the case both for the WT and *mad2*, this potentially indicates an important role of MTs for regulating directly or indirectly interphase duration.

It is noteworthy, that we observed meiocytes, which at the time they were placed on oryzalin, were in pachytene evident by the half-moon structure formed by microtubules around the nucleus (Prusicki et al., 2019). We conclude that, once the commitment to the meiotic program has been made, meiosis progresses even if essential components like microtubules are not present. So, on the one hand, plant cells seem to be able to flexibly silence cell cycle checkpoints such as the SAC or the pachytene checkpoint in order to progress with the meiotic program (this work and De Jaeger-Braet et al., 2021), but on the other hand, cannot escape from their commitment to the program itself.

In the single-cell algae *Chlamydomonas reinhardtii* cells undergo several rounds of DNA replication without nuclear division under microtubule-destabilizing conditions (Tulin & Cross, 2014) and lagging chromosomes as well as cytokinesis failure in the bryophyte *Physcomitrium patens* leads to somatic cell polyploidization (Kozgunova et al., 2019). Similarly, application of microtubule poisons, like colchicine, on mitotic cells of various angiosperms led to the re-formation of a NE without prior chromosome segregation (Molè-Bajer, 1958; Nebel & Ruttle, 1938), suggesting that NE reformation upon severe stress conditions is the rule rather than the exception in the plant lineage. Besides, colchicine and other drugs have long been exploited in plant breeding for generating polyploid offspring (Younis et al., 2014). Considering the prevalence, the challenges but also the opportunities that whole-genome duplication events present (Bomblies et al., 2016; Comai, 2005; Hollister, 2015; Panchy et al., 2016) as well as the evolutionary driving force of hybridization followed by polyploidization (Soltis & Soltis, 2009), we hypothesize that evolution in the plant lineage has been shaped by the weak nature of the SAC. A weak SAC could have facilitated major genomic changes to take place and hence, contribute to plant evolution.

### Mutants in *BUB3.3* exhibits an unexpectedly long metaphase I and synthetic lethality in combination with mutants in other SAC components

Given a differential contribution of several SAC components in regulating metaphase duration in mammals (Meraldi et al., 2004) and yeast (Cairo et al., 2020; Vanoosthuyse et al., 2009; Yang et al., 2015), we next measured the metaphase I duration in the different SAC mutants. To this end, we divided the SAC mutants into two groups: (a) mutants of SAC genes that encode for kinetochore-bound proteins (*mad1* and *mps1*) and (b) mutants of SAC genes that encode for proteins which are putatively components of the MCC (*bmf2* and *bub3.3*) (Komaki & Schnittger, 2017). Additionally, we looked at *mad2* which belongs to both groups.

As expected, all three mutants of SAC components that encode for kinetochore-bound proteins had an ∼ 30% shorter metaphase I duration compared to the WT (Fig. 4A and Video 9). Metaphase I duration in *bmf2* was ∼ 20% shorter compared to the WT (Fig. 4B and Video 10). In striking contrast to all other SAC mutants, in *bub3.3* metaphase I was ∼ 50% longer compared to the WT (Fig. 4B and Video 10). After confirming that *bub3.3* is a null allele, we also observed the prolonged metaphase I duration in a second null allele of *BUB3.3*, *bub3.3-2*, as well as an F1 cross between homozygous *bub3.3* and *bub3.3-2* plants (Fig. EV4B, L, M). Furthermore, metaphase I duration could be rescued to WT levels by the introgression of genomic *BUB3.3* expressed under its own promoter in the *bub3.3* background in three independent lines (Fig. EV4H).

**Figure 4.**
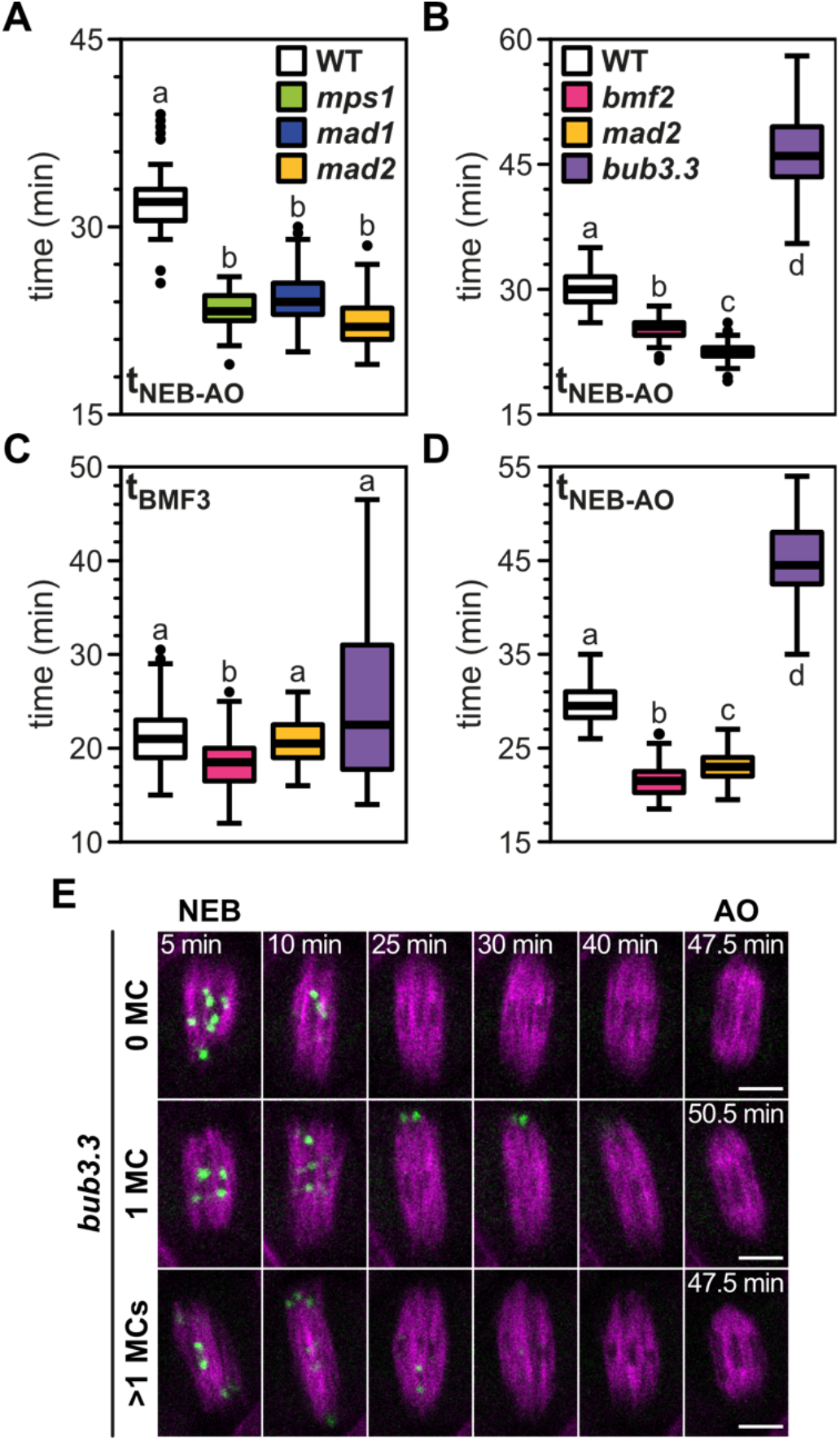
*bub3.3* mutants exhibit prolonged metaphase I duration and delayed homologous chromosome pair congression to the metaphase plate. (A,B) Quantification of metaphase I duration of WT and several SAC mutants whose proteins are either on the kinetochore **(A)** or presumably part of the MCC **(B)**. **(C,D)** Quantification of BMF3 kinetochore signal **(C)** and metaphase I **(D)** duration in the WT and SAC mutants. **(E)** Representative *bub3.3* cells with 0, 1 or multiple mislocalizing chromosome pairs (MC) in metaphase I. BMF3 is fused to GFP (green) and TUA5 to TagRFP (magenta). 2D movies were recorded with a frame rate of 30 seconds. For **(A)**, we quantified 68-98 cells from 11-16 anthers per genotype. For **(B)**, we quantified 67-86 cells from 16-17 anthers per genotype. For **(C,D)**, we quantified 62-154 cells from 10-28 anthers per genotype. Different letters indicate significance (P < 0.01; pairwise comparison using Kruskal-Wallis test for independent samples with Bonferroni correction for multiple tests). Scale bars: 5 µm.

We became particularly interested in *BUB3.3* due to the contradictory role of its homolog Bub3 observed in yeasts. Both in budding and fission yeast, *bub3*-depleted mitotic cells exhibit a prolonged AO delay because of – but not limited to – inefficient APC/C activation (Vanoosthuyse et al., 2009; Yang et al., 2015). Intriguingly, in budding yeast, *bub3*-depleted meiotic cells have a shorter metaphase I and II duration than WT most probably due to a disruption of the phosphorylation-dephosphorylation balance at the kinetochore (Cairo et al., 2020).

Considering that Bub3 has been implicated in proper chromosome segregation both in yeast (Cairo et al., 2020, 2023; Vanoosthuyse et al., 2009; Yang et al., 2015) and human cells (Logarinho et al., 2008), we scored pollen viability in *bub3.3* and found that it was the lowest among the WT, *bmf2* and *mad2* (Fig. EV5A). To start untangling the role of *BUB3.3 in planta*, we, then, generated double mutants between *bub3.3* and mutants in core SAC components. Similar experiments in yeast mitosis have revealed a diversified role of Bub3. In fission yeast, the *mad2* mutation is epistatic, while in budding yeast, the *bub3* mutation is epistatic in the *bub3 mad2* double mutant (Vanoosthuyse et al., 2009; Yang et al., 2015). Hence, we chose to cross *bub3.3* with *mad1*, *mad2* and *bmf2* mutants to test which of the mutations is either epistatic or additive *in planta*. Notably, we could not obtain any viable double homozygous mutant with *bub3.3* (Fig. EV5B). Genotyping the progeny of *bub3.3 −/− mad2 +/−* and *bub3.3 +/− mad2 −/−*, we obtained a ratio of 89:150 WT:Heterozygous seedlings indicating that the double mutant is most probably sporophytic (embryonic) lethal. Consistent with a predominant sporophytic defect, we found that the transmission of the *bub3.3* allele was neither reduced when using the mutant as a male (1:1.48 WT:*bub3.3* T-DNA) nor as a female (1:1.53 WT:*bub3.3* T-DNA) parent in reciprocal crosses between *bub3.3 +/− mad2 −/−* and the WT (Fig. EV5C).

In contrast to double mutants of SAC components with *bub3.3*, we found that the combinations of other SAC mutants did not result in lethality. We could successfully obtain all homozygous double mutant combinations between *mad1*, *mad2* and *bmf2*, i.e., *mad1* x *bmf2*, *mad2* x *bmf2* and *mad1* x *mad2* (Fig. EV5B and Video 11). We then measured metaphase I duration in these double mutants as it has been shown in HeLa cells that double knockdown of BubR1 and Mad2 has an additive effect suggesting that these proteins might inhibit the APC/C through parallel pathways as well (Meraldi et al., 2004). However, this does not seem to be the case for Arabidopsis as no additive effect on metaphase I duration could be observed in any of the double mutant combinations we generated (Fig. EV5D).

### Prolonged metaphase I duration in *bub3.3* correlates with mislocalizing homologous chromosome to the spindle pole

The synthetic lethality of *bub3.3* with other SAC mutants, which has never been observed in any other system, prompted us to hypothesize that BUB3.3 might be predominantly regulating chromosome congression rather than being a core SAC component. In mammals and yeast, Bub3 interacts with both Bub1 and BubR1 and recruits them to the kinetochore via its recognition of and interaction with phosphorylated MELT motifs in Knl1/Spc105 (London et al., 2012; Overlack et al., 2015; Primorac et al., 2013; Shepperd et al., 2012; Vleugel et al., 2013, 2015). However, in plants BMF1 and BMF2, the respective homologs of Bub1 and BubR1, as well as BMF3 have lost the Gle2-binding sites (GLEBS domain), via which they interact with Bub3 (Larsen et al., 2007; Taylor et al., 1998; Wang et al., 2001). Consistently, it was recently shown in maize (*Zea mays*), that ZmBub3 does not interact in a yeast-two-hybrid assay neither with ZmKnl1 nor with ZmBmf1/2/3 (Su et al., 2021). Matching these interaction data, we found that the localization of both BMF1 and BMF3 in *bub3.3* was indistinguishable from the WT in metaphase I (Fig. 4E, EV5E and Video 12, 13).

To approach the question of what the function of BUB3.3 could be, we followed the BMF3 reporter in the *bub3.3* background and compared the phenotype to *bmf2* and *mad2*. In agreement with our previous results (Fig. 4B), the metaphase I duration of *mad2* and *bmf2* were shorter while in *bub3.3* it was longer than in the WT (Fig. 4D and Video 13). In contrast to metaphase I, the BMF3 kinetochore signal duration in *bub3.3* was not consistently longer than in the WT and exhibited a large variation in comparison to the other genotypes (Fig. 4C). We recognized that the *bub3.3* cells fall into two groups, with ∼ 60% of the cells having a similar distribution with the other genotypes and ∼ 40% having a prolonged BMF3 kinetochore signal persistence (Fig. EV5F). Upon closer inspection, we realized that *bub3.3* cells with a prolonged BMF3 kinetochore signal persistence exhibited a characteristic chromosome mislocalization phenotype, with individual or multiple homologous chromosomes localizing to the spindle pole(s) (Fig. 4E). The two paired kinetochores tended to be at an almost right angle to the long axis of the spindle, indicating that most probably these kinetochores were unattached, poorly attached or syntelically attached to spindle microtubules. This would also explain why these kinetochores are persistently highlighted by BMF3 in contrast to kinetochores that aligned efficiently in the metaphase plate. We did not observe a similar defect neither in *bmf2* nor *mad2*, suggesting that it is specific to *bub3.3*. All these observations were confirmed with a BMF1 reporter, introgressed in *bub3.3* as well *bmf2* and *mad2*, that persistently labels the kinetochore independently of kinetochore-microtubule attachment status (Fig. EV5E).

Based on the synthetic lethality of *bub3.3* with all other SAC mutants tested and its unique chromosome mislocalization phenotype, BUB3.3 does not seem to be primarily involved in the plant SAC, but in a parallel pathway.

### *BUB3.3* is not a *bona fide* SAC component

Although, we could observe mislocalizing chromosome pairs in ∼ 40% of *bub3.3* cells, all of them had a comparatively equally long metaphase I (Fig. 4B, D and EV4B, H). So, we hypothesized that in our 2D movies we were likely underestimating the number of cells with aberrant chromosome behavior in *bub3.3*. Thus, we proceeded to generate 3D movies of WT, *bub3.3*, *mad2* and *bmf2* to capture the entire cell volume. Our results regarding BMF3 kinetochore signal and metaphase I duration in 3D movies (Fig. 5A, B, EV4A, B and Video 14, 16) mirrored almost perfectly the results obtained in 2D movies (Fig. 4C, D) confirming the large variation of SAC activity in *bub3.3* cells. Then, we quantified the proportion of cells exhibiting at least one mislocalizing chromosome pair (MC). This was the case for ∼ 75% of *bub3.3* cells compared to ∼ 10% for all other genotypes (Fig. 5B) and the ∼ 40% of *bub3.3* we had obtained from the 2D movies (Fig. EV5F). The MCs in *bub3.3* stayed at the spindle pole from several minutes to over one hour while in the other mutants they averaged out at ∼ 5 minutes (Fig 5B). This reflected fully and essentially explained the large variation in SAC activity in *bub3.3* and reinforced the notion that the lack of proper chromosome-spindle microtubule attachment seems to be a stochastic, cell-autonomous process. Additionally, the distribution of the number of MCs per cell revealed that most *bub3.3* cells had only one mislocalizing pair in metaphase I, however, a substantial amount had two or more, which was rarely – if ever – the case in the WT, *mad2* and *bmf2*. We consistently observed MCs both on the same or opposite spindle poles with no clear preference, suggesting that this process is stochastic. Finally, with the exception of a single missegregation event observed in a *mad2* cell, missegregation events were regularly observed in *bub3.3* meiocytes.

**Figure 5.**
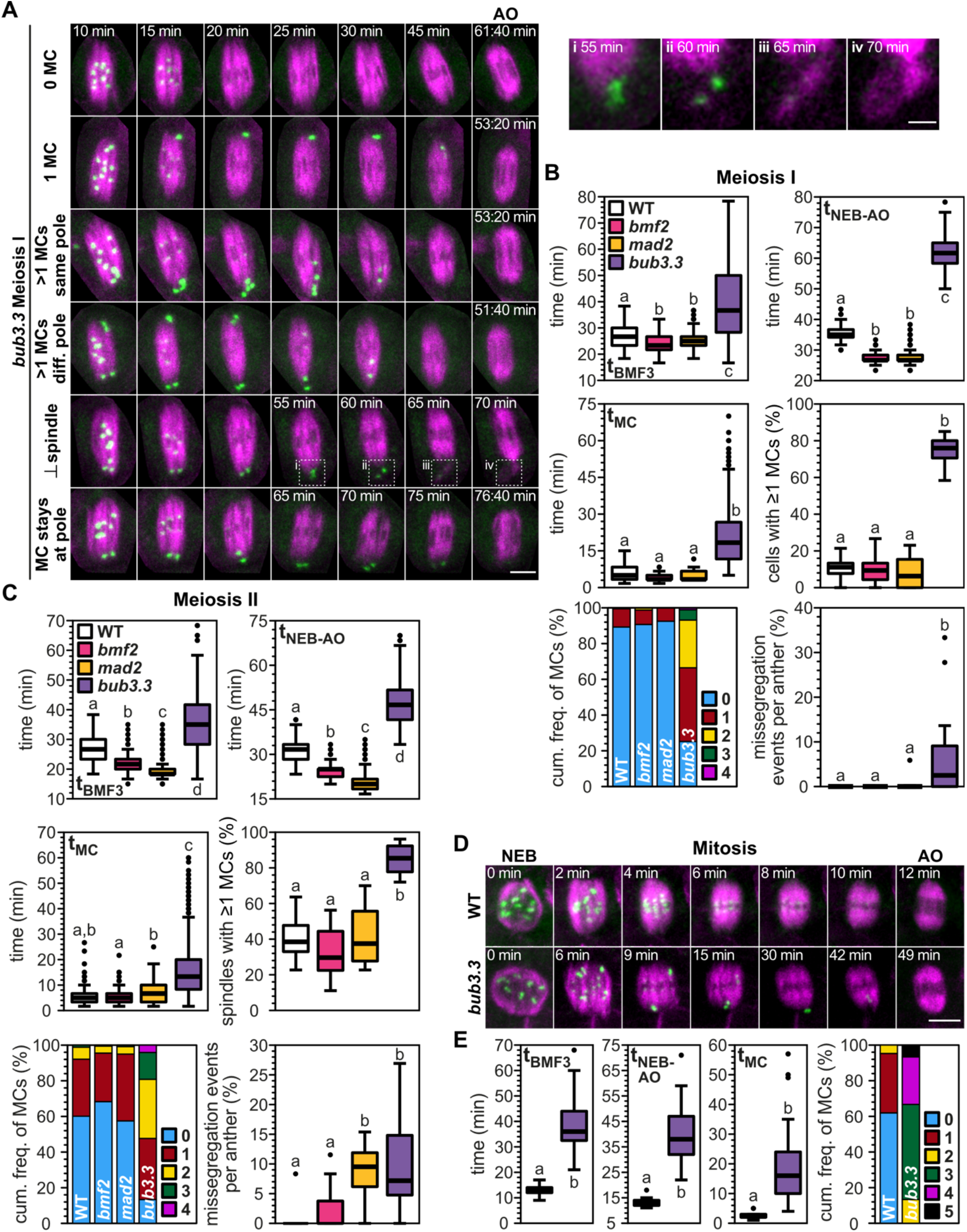
*bub3.3* cells exhibit similar chromosome congression defects in meiosis and mitosis. **(A-C)** We recorded 3D movies of WT, *bmf2*, *mad2* and *bub3.3* anthers in meiosis I **(A,B)** and meiosis II **(C)** and movies of WT and *bub3.*3 in mitosis **(D,E)**. **(A)** Representative maximum projections of *bub3.3* cells with 0, 1 or multiple mislocalizing chromosome pairs (MC) in metaphase I. BMF3 is fused to GFP (green) and TUA5 to TagRFP (magenta). In cells, where chromosome pairs persistently stay at the spindle pole, they missegregate either through the formation of a perpendicular (***⊥***) spindle or by staying at the spindle pole. In both cases the BMF3 signal slowly fades indicating a silencing of the SAC. **(i-iv)** Insets highlight the formation of a perpendicular spindle. **(B,C,E)** We quantified the duration of the BMF3 signal on kinetochores, duration of metaphase, duration of chromosome pairs mislocalizing to the pole(s), percentage of cells per anther with at least one MC, the cumulative frequency of the maximum number of MCs per cell and the percentage of missegregation events per anther. **(D)** Representative maximum projections of WT and *bub3.3* cells. 3D movies were recorded with a frame rate of 100 seconds **(A-C)** or 1 minute **(D,E)**. For **(B)**, we quantified 195-263 cells from 13-18 anthers per genotype in meiosis I. For **(C)**, we quantified 341-459 spindles from 15-22 anthers per genotype. Different letters indicate significance (P < 0.01; pairwise comparison using Kruskal-Wallis test for independent samples with Bonferroni correction for multiple tests). For **(E)**, we quantified 15-21 cells per genotype. Different letters indicate significance (P < 0.01; Mann-Whitney U test for independent samples). Scale bars: 5 µm **(A)** and 2 µm **(i-iv)**.

All defects observed in *bub3.3* in metaphase I were recapitulated in *bub3.3-2* and the F1 cross between homozygous *bub3.3* and *bub3.3-2* plants in 3D movies (Fig. EV4A-F and Video 14). Moreover, in 3D movies of metaphase I in the three rescue lines, the genomic *BUB3.3* construct brought the quantified parameters back to WT levels corroborating that loss of *BUB3.3* and not any other gene causes the above-described chromosome segregation defects (Fig. EV4G-K and Video 15).

Similar results were obtained for metaphase II (Fig 5C and Video 17). Here, we analyzed the two spindles in each cell individually, because (1) it was not always possible to capture both spindles due to their angle relative to our imaging axis and (2) in cells where we could image both spindles, we did not find any apparent correlation between the behavior of the two spindles with regard to MCs. In metaphase II, we observed for all genotypes a higher proportion of spindles with at least one MC compared to metaphase I. A similar trend was observed for the missegregation events per cell, which were more abundant compared to metaphase I.

Having analyzed the *bub3.3* phenotype in metaphase I and II, we asked whether this phenotype is also found in mitosis. This is especially interesting considering that partially opposing phenotypes in mitotic and meiotic *bub3*-depleted cells were described in budding yeast (Cairo et al., 2020; Yang et al., 2015). For this, we recorded 3D movies of mitotic root epidermal cells in the WT and *bub3.3* background. Recapitulating essentially our observations in meiosis, *bub3.3* cells stayed much longer in metaphase than the WT, exhibiting a prolonged BMF3 kinetochore signal due to MCs which persisted for up to ∼ 1 hour (Fig. 5D, E and Video 18). Additionally, in all observed *bub3.3* cells, we found at least two MCs. Given the progressive worsening of the *bub3.3* phenotype from metaphase I through metaphase II over to mitosis, it is tempting to speculate that perhaps kinetochore size, which becomes progressively smaller in these divisions, might play a role in that. It would be an interesting line of research to follow in the future, for instance in mutants with reduced centromere size (Capitao et al., 2021), considering similar work in animal cells (Drpic et al., 2018).

## Conclusion

Surveillance of faithful chromosome segregation in meiosis is of key importance for fertility and transgenerational genome stability. Thus, studying the SAC is of great relevance to understand not only fundamental aspects of plant reproduction and evolution but is also highly relevant for plant breeding. Importantly, male meiocytes, as demonstrated in this study, are a powerful model system to also study general aspects of the SAC due to their high level of synchronicity and their accessibility through an established live-cell imaging approach (Prusicki et al., 2019) together with the possibility to apply drugs such as the MT poison oryzalin (Sofroni et al., 2020; this work). Here, we have quantitatively assessed key parameters and properties of the SAC, such as the timing of its assembly and disassembly, setting the foundation for future studies of this checkpoint (Fig. 6). Our work has also allowed us to assign BUB3.3 a novel role in kinetochore-microtubule attachment and chromosome congression instead of a function in the SAC (Fig. 6). These observations led us to a refined and temporally resolved model of the composition and regulation of the Arabidopsis core SAC comprising AUR1/2/3, BMF1, BMF2, BMF3, MAD1, MAD2 and MPS1 (Fig. 6).

**Figure 6.**
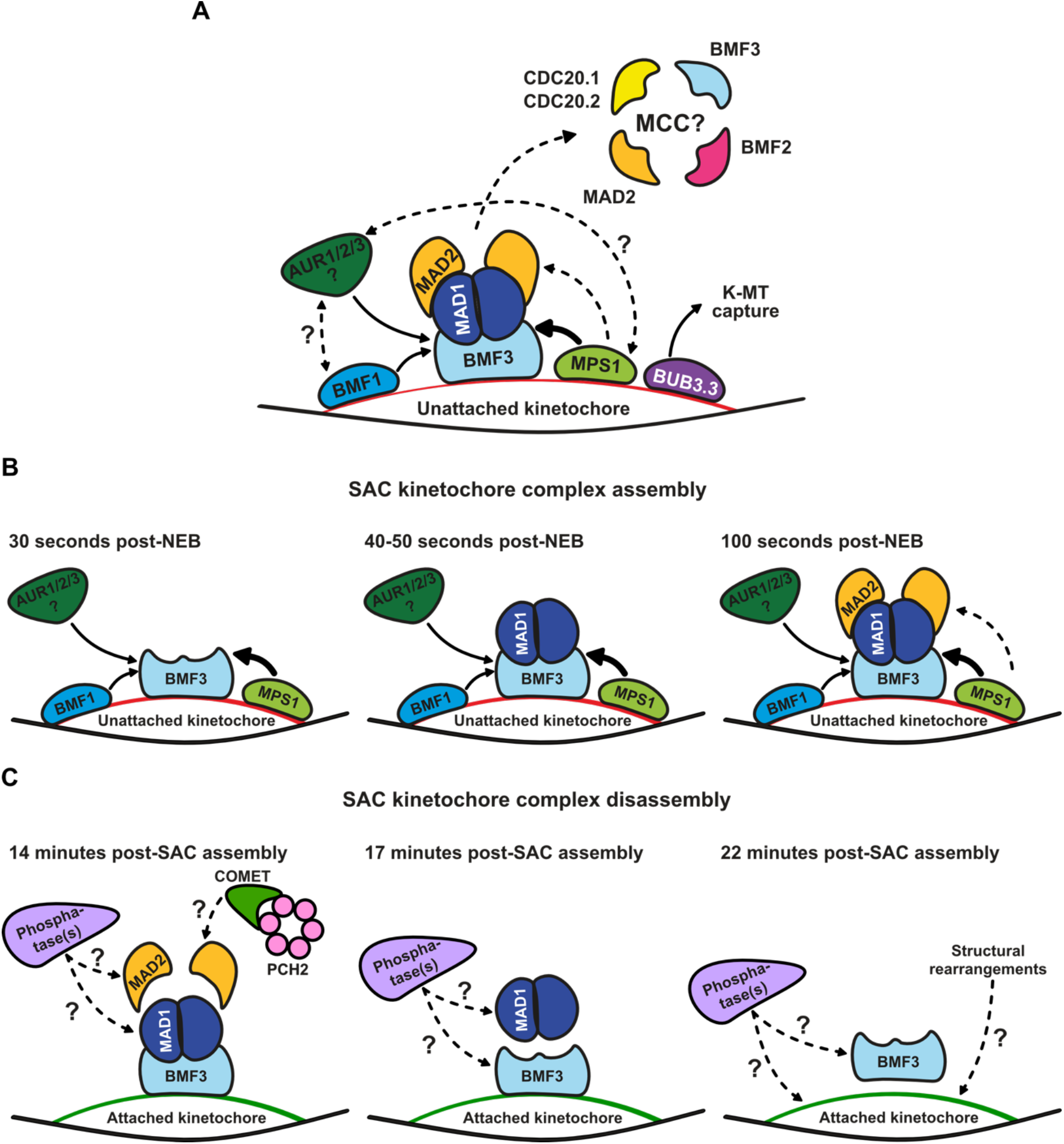
A revised model of the Arabidopsis SAC and the role of BUB3.3. Solid line arrows indicate conclusions from this study, dotted line arrows indicate conclusions from Komaki & Schnittger (2017), while dotted line arrows with question marks indicate hypotheses. **(A)** The timing as well as the protein amount of BMF3 recruited to the kinetochore is influenced by AURORA kinases, BMF1, and MPS1. Most probably, AUR3 – and not AUR1/2 – is responsible for the timely and sufficient recruitment of BMF3 to the kinetochore as it is localized at the kinetochore during metaphase. It remains to be determined whether AURORA kinase(s), BMF1, and MPS1 cross-regulate each other like in mammalian cells. Finally, BUB3.3 is probably not a *bona fide* SAC component and this is why it is probably not a component of the MCC in plants. The composition of the plant MCC still remains to be resolved. Additionally, BUB3.3 is not mediating the kinetochore anchorage of BMF1 or BMF3 in Arabidopsis. Thus, we hypothesize that similarly to other organisms BUB3.3 sits at the outer kinetochore, but has evolved novel functions. BUB3.3 is probably involved in kinetochore-microtubule (K-MT) attachment formation and chromosome congression. Conclusively, *BUB3.3* appears to have evolved a novel role in plants and, in contrast to other organisms, to not be a SAC component. **(B)** BMF3, MAD1 and MAD2 are recruited to the kinetochore in a stepwise fashion, where BMF3 comes first, followed by MAD1, and finally, MAD2. According to our data, this process is most probably regulated either directly or indirectly through kinases. **(C)** The disassembly of the complex happens in the opposite order. Phosphatases have been shown to counteract the action of kinases and promote SAC silencing (Lara-Gonzalez et al., 2021). Their combined action fine-tunes SAC activity and eventually silences the SAC. The PCH2-COMET complex which is present in Arabidopsis (Balboni et al., 2020) might be involved in removing MAD2 from the complex, since it has been shown to be involved in SAC silencing in mammalian cells (Lara-Gonzalez et al., 2021). Finally, outer kinetochore components undergo structural rearrangements upon K-MT attachment formation (Magidson et al., 2016; Roscioli et al., 2020), which might play a role in the disassociation of SAC complex components from the outer kinetochore.

## Materials and Methods

### Plant materials

We used the *Arabidopsis thaliana* (Arabidopsis) accession Columbia-0 (Col-0) as the wild type (WT) throughout this research and all mutant lines are in the Col-0 background. We used the following T-DNA insertion lines, which we obtained from the Nottingham Arabidopsis Stock Center (NASC) (http://arabidopsis.info/): SALK_122554 (*bmf1*), SAIL_303_E05 (*bmf2*), GABI_084G06 (*bmf3*), SALK_041372 (*bub3.3*), SALK_022904 (*bub3.3-2*), SALK_073889 (*mad1*), SAIL_191G06 (*mad2*), GABI_663H07 (*mps1*). All lines, except for *bub3.3-2*, were previously used in Komaki & Schnittger (2017). For *bub3.3-2*, we used the same genotyping primers as for *bub3.3* (Komaki & Schnittger, 2017).

For mitosis imaging, seedlings were grown for 4 days on a solid medium containing half-strength Murashige and Skoog (MS) medium, 1% (w/v) sucrose, 0.8% (w/v) agar and pH 5.8 in a growth chamber (16h light at 21 °C/8h dark at 18 °C). For imaging and phenotypic analyses of meiosis, 7- to 10-day old seedlings grown on plate as described above were transferred to soil and grown for 5-6 weeks in growth chambers (16h light at 21 °C/8h dark at 18 °C).

### Genotyping, T-DNA mapping and RT-PCR

Primers for genotyping of T-DNA insertion lines are described in Komaki & Schnittger (2017). For, *bub3.3-2* we used the same genotyping primers as for *bub3.3* (Komaki & Schnittger, 2017). To map the T-DNA insertion in *bub3.3* and *bub3.3-2*, we used the left border primer for the T-DNA SALK_Lb1.3 (ATTTTGCCGATTTCGGAAC). For the <u>r</u>everse transcription-<u>p</u>olymerase <u>c</u>hain <u>r</u>eaction (RT-PCR), we isolated RNA from ∼ 200 9-day old seedlings per genotype grown as described above. The seedlings were collected in an Eppendorf tube, flash frozen in liquid nitrogen and RNA was isolated according to the manual of the RNA-isolation kit (innuPREP Plant RNA Kit, Analytik Jena AG). cDNA synthesis was performed according to the manual of the cDNA synthesis kit (Thermo Scientific RevertAid Reverse Transcriptase, Thermo Fisher Scientific) using the Oligo(dT)18 primer. According to The Arabidopsis Information Resource (TAIR, https://www.arabidopsis.org/), there are two splice variants of *BUB3.3*, a long (Splice variant 1) and a short (Splice variant 2) variant. We performed two PCR reactions for each genotype using primers TCGGAATTCGAAATTGGGGA (fwd) and TGGTTGACACGAGTCAGTTCT (rev) to amplify the full-length of Splice variant 1 and primers TCTCCTCGTTGCTTCTTGGG (fwd) and GGCCAAGAGTTCTCCAGTGT (rev) to amplify both splice variants simultaneously. We used histone H2A as a positive control and primers ATGGCGGGTCGTGGTAAAACACTCGGATCT (fwd) and TCAATCGTCTTCAGCAGATGGCTTGGAAGCACC (rev).

### Plasmid construction and line generation

All constructs – except for *BUB3.3::gBUB3.3* – were previously generated in Komaki & Schnittger (2017). To generate the *BUB3.3::gBUB3.3*, construct the *BUB3.3* gene, including 2 kb upstream of the start codon and 0.5 kb downstream of the stop codon, was amplified by PCR and cloned into *pDONR221*, followed by LR recombination reaction with the destination vector *pGWB501*. We used the following primers: <u>GGGGACAAGTTTGTACAAAAAAGCAGGCT</u>GCTCTAGATTTGCTTGTT (forward) and <u>GGGGACCACTTTGTACAAGAAAGCTGGGT</u>GATTCAGCGAAATCGGAA (reverse). The underlined part of the sequence indicates the *attB*-sites.

We introgressed all reporters from the WT into the mutant backgrounds through genetic crossing. We wanted to ensure that all lines carry the same reporter insertion(s), thus eliminating the possibility that variation in expression level due to insertion site would influence our observations.

### Pollen viability

We used the Peterson staining protocol to score pollen viability (Peterson et al., 2010). Three mature flowers with dehisced anthers were dipped in 20 µl Peterson staining solution (10% ethanol, 0.01% malachite green, 25% glycerol, 0.05% fuchsin, 0.005% orange G, and 4% glacial acetic acid) and incubated for 1h at 95 °C. The samples were analyzed the same day.

### Live-cell imaging

Live-cell imaging of anthers was performed using the protocol described in Prusicki et al. (2019) with slight variations. 40-80 flower buds were detached from the inflorescence stem keeping the pedicel intact and were placed on live-cell imaging medium as described in Prusicki et al. (2019). For 2D live-cell imaging, samples were prepared either the same day or the day before and incubated overnight in the dark. For 3D live-cell imaging, samples were prepared the day before and incubated overnight in the dark.

2D live-cell imaging of anthers was performed on a Zeiss LSM780 upright confocal laser scanning microscope (CLSM) equipped with a GaAsP-detector and a W-plan-Apochromat 40x/1.0 DIC M27 or a W-plan-Apochromat 63x/1.0 DIC water-dipping lens. 8-bit images were acquired using bi-directional scanning every 5 seconds to 2 minutes with 100-120 nm lateral pixel resolution, 4 times line average, 1.3-2.7 µs pixel dwell time and a pinhole size of 1-2 AU (for the GFP detection channel). GFP was excited at 488 nm and emission was detected at 489-550 nm. TagRFP was excited at 561 nm and emission was detected at 565-650 nm. GFP and TagRFP were detected either sequentially using line-scan mode or simultaneously to increase the frame rate. In the latter case, to prevent crosstalk, TagRFP emission was detected at 576-655 nm.

3D live-cell imaging of anthers was performed on a Zeiss LSM880 upright CLSM equipped with a GaAsP-detector and a W-plan-Apochromat 40x/1.0 DIC M27 water-dipping lens. 16-bit images were acquired using bi-directional scanning every 100 seconds to 2 minutes with 84-96 nm lateral pixel resolution, 2 times line average, 1.04-1.18 µs pixel dwell time and a pinhole size of 1.51 AU (for the GFP detection channel). z-stacks were acquired with a step-size of 0.93 µm. GFP and TagRFP were excited at 488 nm and emission was detected simultaneously at 490-553 nm and 579-651 nm, respectively.

For imaging of mitosis, we used 4-day old seedlings grown vertically as described above. Before imaging, the seedlings were carefully lifted from the growth medium and placed in a drop of liquid half-strength MS-medium on a glass-bottom dish. 5-8 seedlings were placed on each dish and then, we covered the roots with a strip of solid half-strength MS-medium.

3D live-cell imaging of mitotic epidermal root cells was performed on a Leica TCS SP8 inverted CLSM equipped with three hybrid photodetectors (Leica HyD™) and a HC PL APO 63x/1.2 W motCORR CS2 water-immersion objective. 12-bit images were acquired using bi-directional scanning every 1 min with 99 nm lateral pixel resolution, 4 times line average, at a speed of 700 Hz and a pinhole size of 1.5 AU (for 510 nm emission wavelength). z-stacks were acquired with a step-size of 0.52 µm. GFP and TagRFP were excited at 488 nm and emission was detected simultaneously at 493-555 nm and 575-655 nm, respectively.

### Oryzalin and hesperadin treatment

For the oryzalin treatment, we followed the protocol described in Sofroni et al. (2020) with slight variations. A 10 mM oryzalin (Duchefa Biochemie) stock solution was prepared in dimethyl sulfoxide (DMSO) and stored at −20 °C. 1:10,000 (v/v) stock solution or the equivalent amount of DMSO was added to the medium used for live-cell imaging. We always used freshly prepared plates. Flowers were incubated minimum 4-5 hours in the medium prior to live-cell imaging.

For the hesperadin treatment, we prepared a 10 mM hesperadin (Sigma-Aldrich) stock solution in DMSO and stored it at −20 °C. The appropriate amount of the stock solution was added to the live-cell imaging medium. As a mock treatment, we added 1:1000 (v/v) DMSO to the medium. We always used freshly prepared plates. Flowers were incubated approximately 12 hours in the medium prior to live-cell imaging.

### Post-processing of movies and measurement of fluorescence intensity

We performed all post-processing operations and fluorescence intensity (FI) quantifications of our movies in Fiji (Schindelin et al., 2012).

For 2D movies, we only adjusted the brightness and contrast settings and extracted single cells manually using the rectangle selection tool.

For 3D movies, we performed two different protocols of post-processing for the BMF3 and the TUA5 channel, due to the different nature of the signal (spheres vs complex 3D structure of the spindle). For the BMF3 channel, we used the command “Gaussian blur 3D” with a sigma radius of 1 for x, y and z. For the TUA5 channel, we followed a variation of the protocol presented in Krüger (2017). We first duplicated the movie using the command “Duplicate” with the setting “Duplicate hyperstack”. We renamed the two stacks as “GB_15” and “GB_0.5”. Next, we used the command “Gaussian blur 3D” with a sigma radius of 15 for x, y and z on “GB_15” and with a sigma radius of 0.5 for x, y and z on “GB_0.5”. Using the command “Image calculator” we subtracted “GB_0.5” from “GB_15”. This operation removes the sharper signal from the blurred background and generates the image “Background”. Then, using again the command “Image calculator” we subtracted “Background” from “GB_0.5”, resulting in the image “TUA5_final”. After removing the background noise through these mathematical operations, we used the command “Gaussian blur 3D” with a sigma radius of 0.5 for all dimensions on “TUA5 final” to smoothen the image. Upon post-processing the two channels, we extracted single cells manually using the polygon selection tool, the “duplicate” command for the appropriate z-slice range for each cell and finally the “clear outside” command for each z-slice. Finally, we applied the “Z Project” command with the setting “Max Intensity” for each cell and channel and adjusted the brightness and contrast settings.

To measure the maximum BMF3 reporter FI, we manually drew an ellipse using the elliptical selection tool in a way that all kinetochores labeled with BMF3 were within the ellipse’s boundaries throughout the movie. The FI was measured in all frames from NEB to AO and normalized to the frame of the NEB, which we designated as t_0_. All other timepoints were designated as t_n_, where n is the number of the frame post-NEB. We used the following equation for normalizing the signal intensity: [FI(t_n_) - FI(t_0_)] / FI(t_0_).

### Statistics and figure design

All graphs and statistical operations were generated in IBM SPSS Statistics Version 28 (IBM Corporation).

The boxplots highlight the 25^th^ and 75^th^ percentile indicated by the lower and upper border of the box, respectively, the median with a thick line and the outliers within 1.5 times the interquartile range with the whiskers. Outliers outside this range are indicated by dots above and below the whiskers.

We performed exclusively non-parametric statistical tests and our significance threshold was always P < 0.01. For comparisons between two genotypes, we performed a Mann-Whitney U test. For comparisons between multiple genotypes and/or treatments, we performed pairwise comparisons using a Kruskal-Wallis test for independent samples with Bonferroni correction for multiple tests.

The figures were assembled in Affinity Designer v. 1.10 (Serif).

## Acknowledgments

We thank Dr. Mariana Romeiro Motta (Laboratoire de Reproduction et Développement des Plantes, Lyon, France) for the critical reading of the manuscript. This work was supported by a Ph.D. fellowship to F.B. from the Friedrich-Ebert-Stiftung and core funding of the University of Hamburg (A.S.). Further support was provided by the Ministry of Education, Culture, Sports, Science, and Technology (KAKENHI Grant-in-Aid for Scientific Research on Innovative Areas no. JP19H04864 to S.K.), and Takeda Science Foundation (S.K.).

---

**Figure EV1.**
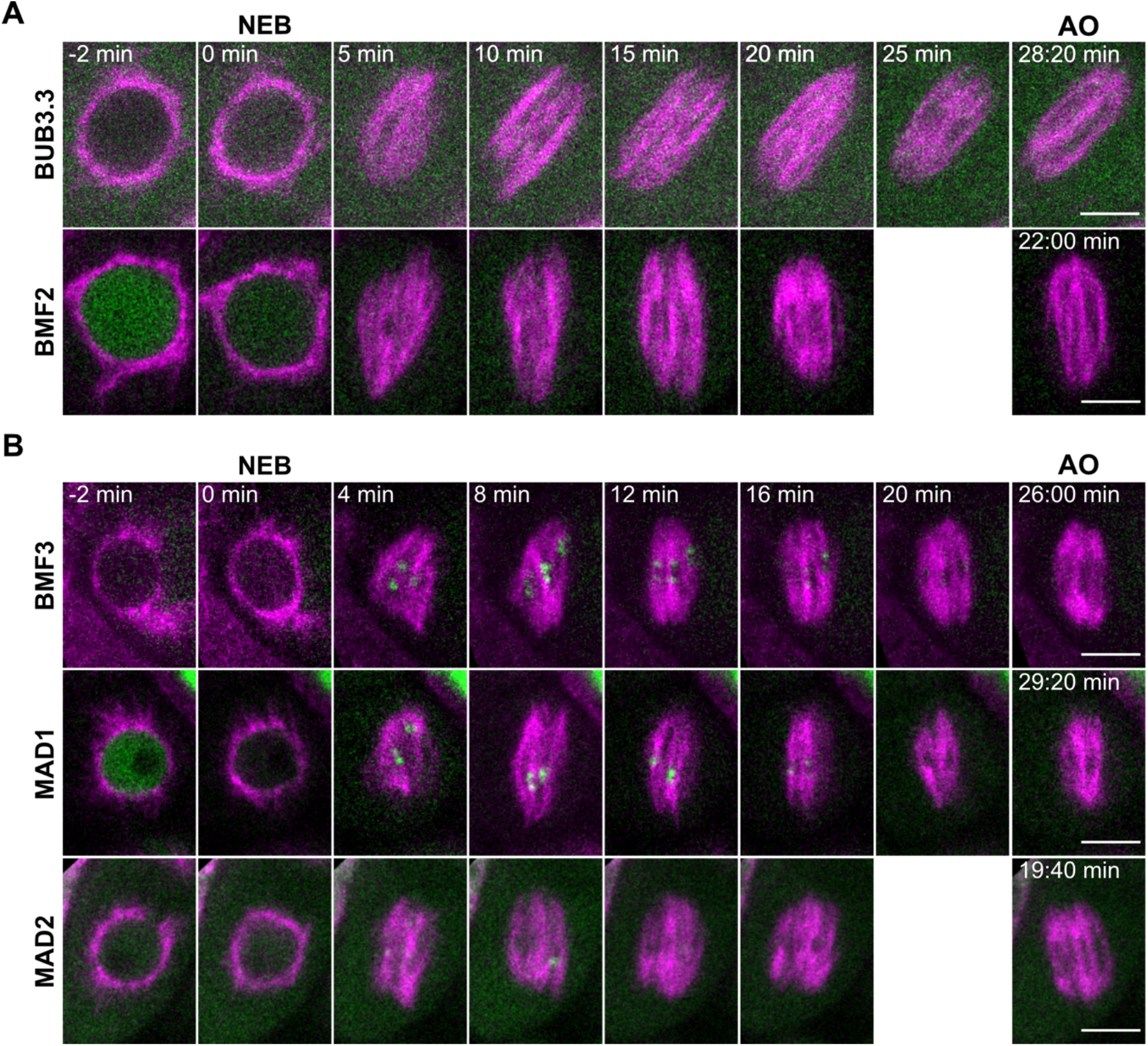
Spatiotemporal localization of functional reporters of core SAC components in meiosis I and II. **(A)** Localization of BMF2 and BUB3.3 fused to GFP together with TUA5 labeled with TagRFP (magenta) in metaphase I. **(B)** Localization of BMF3, MAD1 and MAD2 fused to GFP (green) together with TUA5 labeled with TagRFP (magenta) in metaphase II. 2D movies were recorded with a frame rate of 20 seconds. Scale bars: 5 µm.

**Figure EV2.**
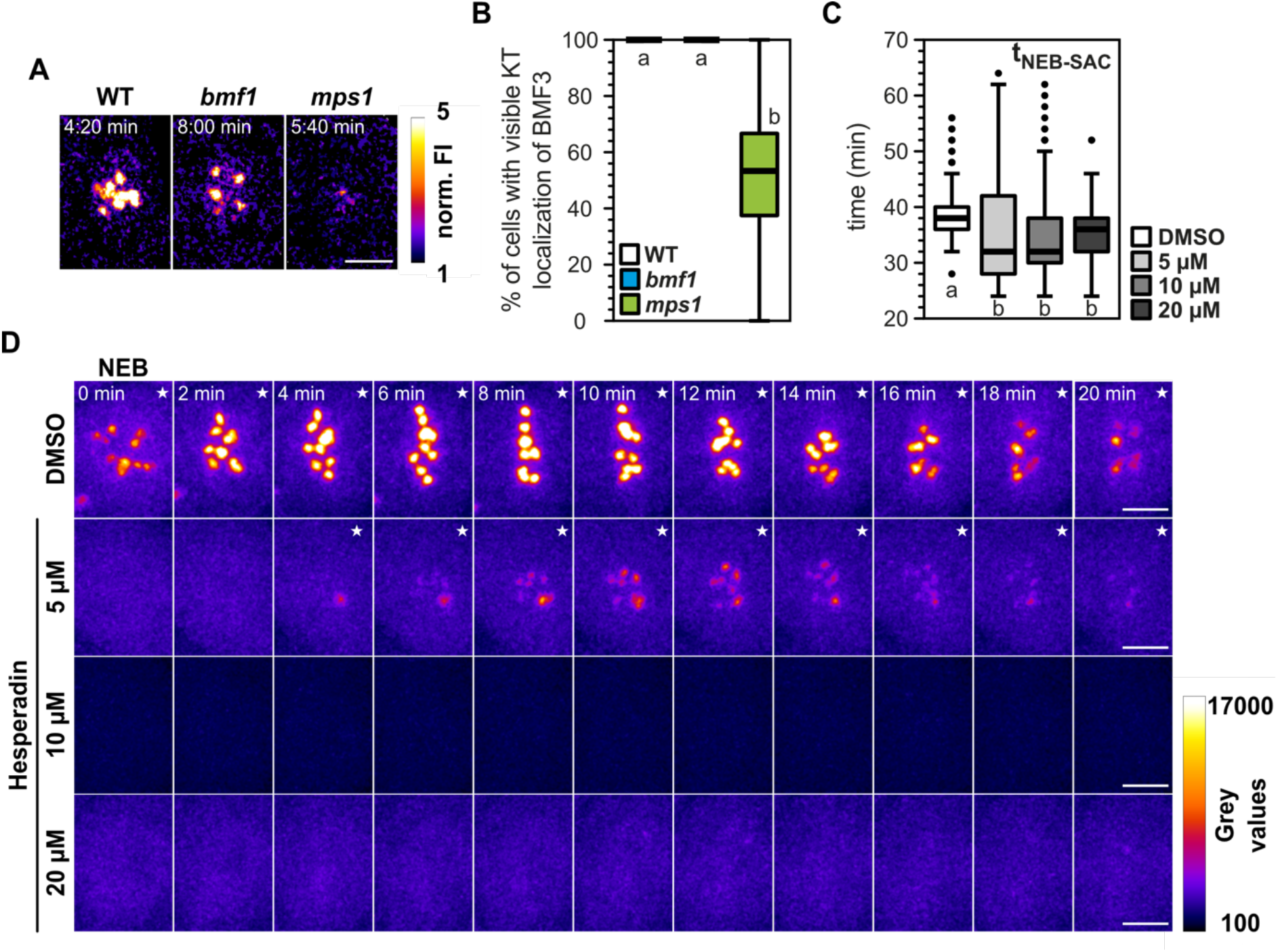
BMF1, MPS1 and AURORA kinases are regulating the strength and timing of the BMF3 recruitment to the kinetochore in metaphase I. **(A)** Timepoints with maximum BMF3 fluorescence intensity (FI) from the same representative cells as in figure 2A. The FI was normalized to the timepoint of NEB. BMF3 is pseudocolored to visualize the decrease in normalized BMF3 signal strength in the *bmf1* and *mps1* backgrounds relative to the WT. **(B)** Percentage of cells per anther with visible BMF3 kinetochore signal during metaphase I. We quantified 75-225 cells from 6-19 different anthers per genotype. **(C)** Metaphase I duration of cells treated with 5, 10 or 20 µM hesperadin or the equivalent amount of DMSO. We quantified 263-339 cells from 13-16 different anthers per treatment. **(D)** Maximum projections of the BMF3 channel of the WT cells shown in figure 2D which were treated with 5, 10 or 20 µM hesperadin or the equivalent amount of DMSO. BMF3 is pseudocolored to visualize the negative correlation between the BMF3 signal strength and hesperadin concentration. The white star (★) on the upper right corner of the frames indicates BMF3 kinetochore localization. Different letters indicate significance (P < 0.01; pairwise comparison using Kruskal-Wallis test for independent samples with Bonferroni correction for multiple tests). Scale bars: 5 µm.

**Figure EV3.**
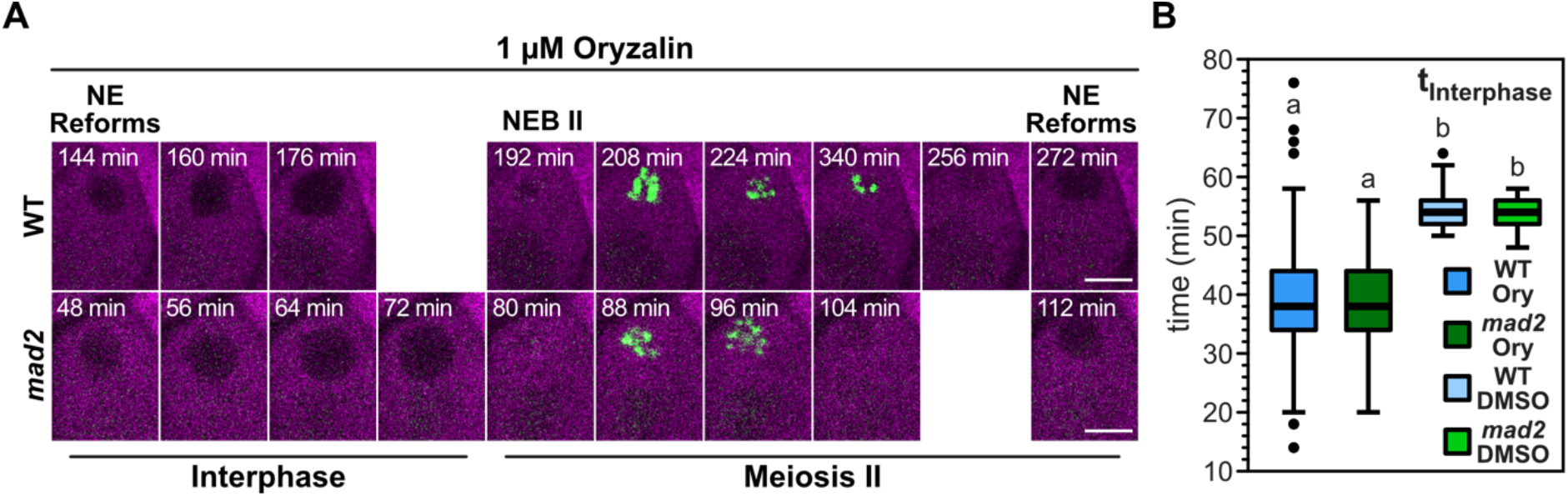
Meiotic cells with depolymerized MTs proceed with meiosis II ending with a second NE reformation. **(A)** Interphase and meiosis II of the same representative WT and *mad2* cells as in Fig. 3C when treated with 1 µM oryzalin. BMF3 is fused to GFP (green) and TUA5 to TagRFP (magenta). **(B)** Quantification of interphase duration in the WT and *mad2*. 2D movies were recorded with a frame rate of 2 minutes. We quantified 69-119 cells from 13-22 anthers per treatment and genotype. Different letters indicate significance (P < 0.01; pairwise comparison using Kruskal-Wallis test for independent samples with Bonferroni correction for multiple tests). Scale bars: 5 µm.

**Figure EV4.**
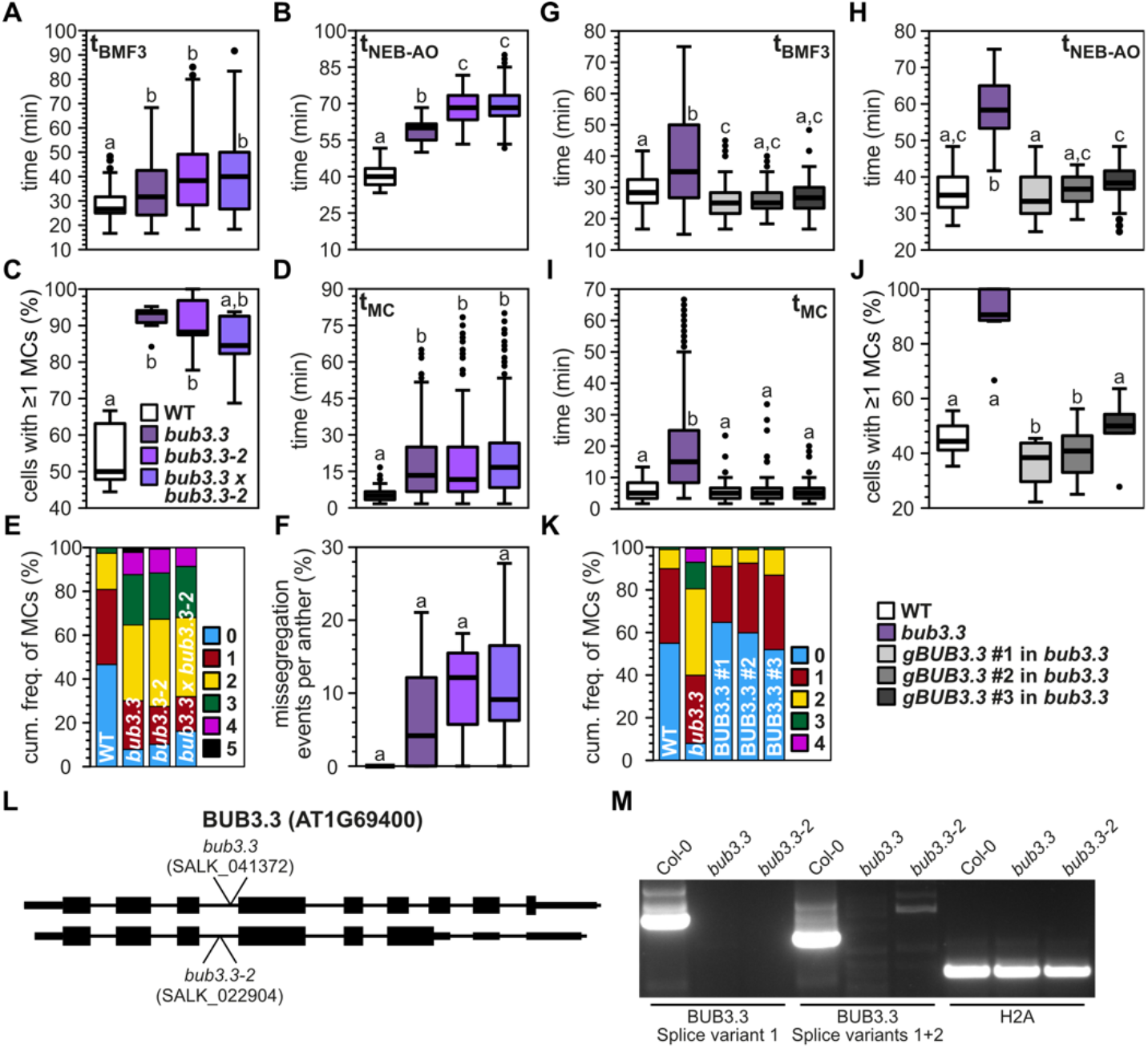
gBUB3.3 rescues the aberrant meiotic phenotype of *bub3.3* in metaphase I, which is further confirmed by a second *bub3.3* allele. (A-F) Both *bub3.3-2* and the double heterozygous F1 cross *bub3.3* +/− x *bub3.3-2* +/− exhibit identical phenotypical defects to *bub3.3*. We quantified the duration of the BMF3 signal on kinetochores **(A)**, duration of metaphase I **(B)**, duration of chromosome pairs mislocalizing to the pole(s) **(C)**, percentage of cells per anther with at least one MC **(D)**, the cumulative frequency of the maximum number of MCs per cell **(E)** and the percentage of missegregation events per anther **(F)**. **(G-K)** Three independent *BUB3.3::gBUB3.3* insertion lines in the *bub3.3* background were genetically crossed with the line expressing BMF3 and TUA5 in the *bub3.3* background. We quantified the duration of the BMF3 signal on kinetochores **(G)**, duration of metaphase I **(H)**, duration of chromosome pairs mislocalizing to the pole(s) **(I)**, percentage of cells per anther with at least one MC **(J)** and the cumulative frequency of the maximum number of MCs per cell **(K)**. 3D movies were recorded with a frame rate of 100 seconds. **(L)** Gene model of the two splice variants of *BUB3.3*. The arrows indicate the left border insertion site of the respective T-DNA. **(M)** RT-PCR of the full length cDNA for either the large or both splice variants. We used *H2A* as a positive control. For **(A-F)**, we quantified 114-128 cells from 6-8 anthers per genotype. For **(G-K)**, we quantified 109-138 cells from 6-8 anthers per genotype. Different letters indicate significance (P < 0.01; pairwise comparison using Kruskal-Wallis test for independent samples with Bonferroni correction for multiple tests).

**Figure EV5.**
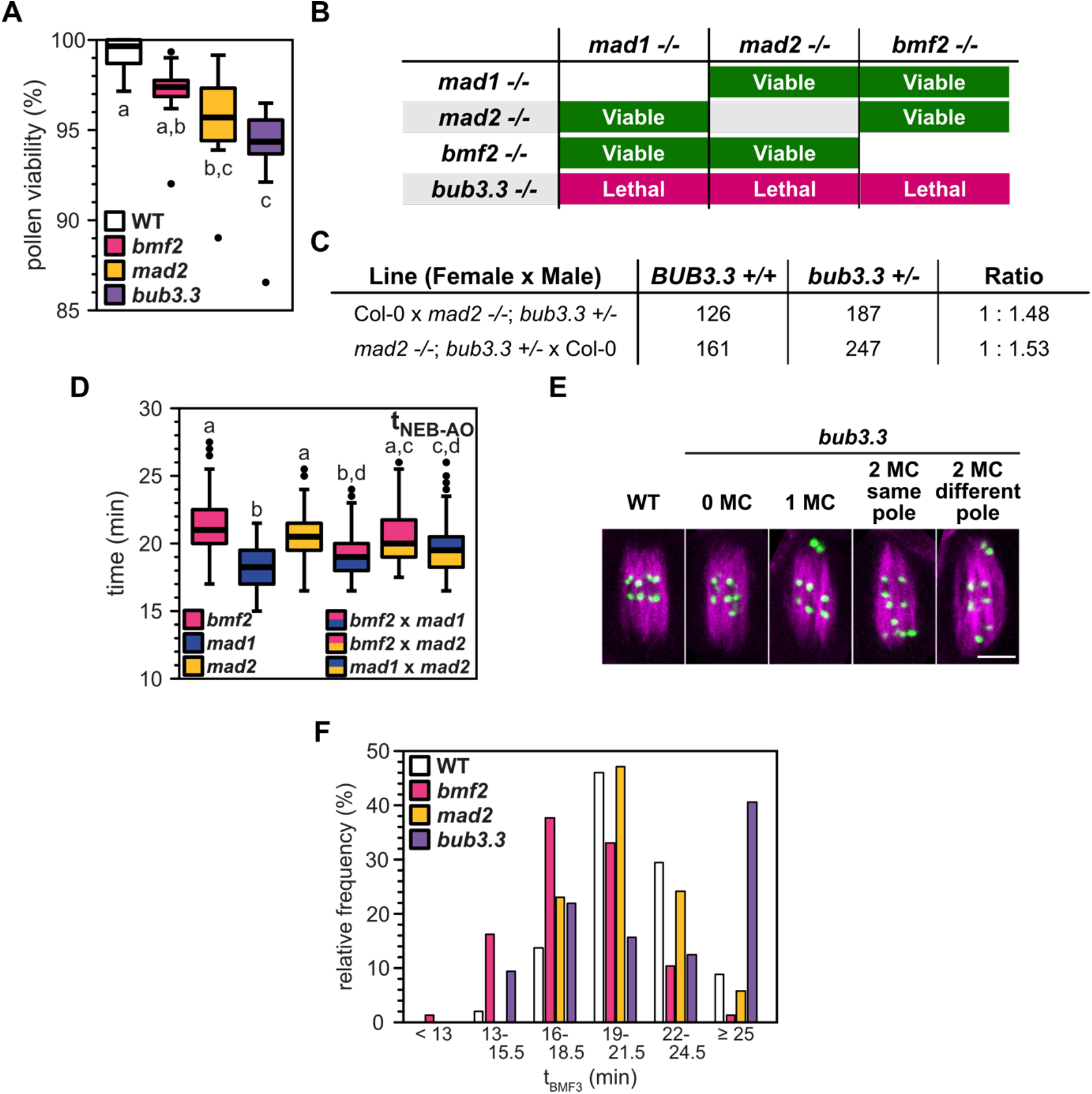
Phenotypic description of SAC single and double mutants. **(A)** Pollen viability of the WT and several SAC mutants. **(B)** Summary of successful and failed double mutant combinations. **(C)** Genotyping results of the F1 progeny of reciprocal crosses between the WT (Col-0) and *mad2 −/−*; *bub3.3 +/−*. **(D)** Quantification of metaphase I duration of viable double mutants. We introgressed the TagRFP::TUA5 reporter in the double mutants through crossing. 3D movies were recorded with a frame rate of 30 seconds. We quantified 79-120 cells from 13-22 anthers per genotype. Different letters indicate significance (P < 0.01; pairwise comparison using Kruskal-Wallis test for independent samples with Bonferroni correction for multiple tests). **(E)** Representative images of WT and *bub3.3* metaphase I. BMF1 is fused to GFP (green) and TUA5 to TagRFP (magenta). From left to right: WT, *bub3.3* with 0 MCs, *bub3.3* with 1 MC, *bub3.3* with 2 MCs on the same spindle pole and *bub3.3* with 2 MCs on opposite spindle poles. Scale bar: 5 µm. **(F)** Relative frequency of the BMF3 kinetochore signal duration in the different groups for each genotype. We quantified the same 64-154 cells from 10-28 anthers per genotype as in Fig. 4C, D.

**Movie EV1 Spatiotemporal analysis of functional SAC reporters with kinetochore localization in metaphase I.** Videos of the representative male meiocytes shown in Fig. 1A. The respective SAC reporter is fused to GFP (green) and TUA5 is fused to TagRFP (magenta) decorating microtubules in metaphase I. The reporters we analyzed are from left to right: BMF3, MAD1, MAD2, MPS1 and BMF1. Movies were recorded with a frame rate of 20 seconds. The playback speed is 10 frames per second.

**Movie EV2 Spatiotemporal analysis of functional SAC reporters with cytosolic localization in metaphase I.** Videos of the representative male meiocytes shown in Fig. S1A. The respective SAC reporter is fused to GFP (green) and TUA5 is fused to TagRFP (magenta). The reporters we analyzed are from left to right: BUB3.3 and BMF2. 2D movies were recorded with a frame rate of 20 seconds. The playback speed is 10 frames per second.

**Movie EV3 Spatiotemporal analysis of functional SAC reporters with kinetochore localization in metaphase II.** Videos of the representative male meiocytes shown in Fig. S1B. The respective SAC reporter is fused to GFP (green) and TUA5 is fused to TagRFP (magenta). The reporters we analyzed are from left to right: BMF3, MAD1 and MAD2. 2D movies were recorded with a frame rate of 20 seconds. The playback speed is 10 frames per second.

**Movie EV4 BMF1 and MPS1 are regulating the timely and sufficient recruitment of BMF3 to the kinetochore in metaphase I.** Videos of the representative male meiocytes shown in Fig. 2A. BMF3 is fused to GFP (green) and TUA5 to TagRFP (magenta). The genotypes we imaged are from left to right: WT, *bmf1* and *mps1*. 2D movies were recorded with a frame rate of 20 seconds. The playback speed is 10 frames per second.

**Movie EV5 AURORA kinases are regulating the timely and sufficient recruitment of BMF3 to the kinetochore in metaphase I.** Videos of maximum projections of the representative male meiocytes shown in Fig. 2D. BMF3 is fused to GFP (green) and TUA5 to TagRFP (magenta). The treatments we performed are from left to right: DMSO (mock), 5 µM hesperadin, 10 µM hesperadin and 20 µM hesperadin. 3D movies were recorded with a frame rate of 2 minutes. The playback speed is 3 frames per second.

**Movie EV6 Metaphase I in a representative WT and *mad2* cell.** Videos of the representative male meiocytes shown in Fig. 3A. BMF3 is fused to GFP (green) and TUA5 to TagRFP (magenta). The genotypes we imaged are from left to right: WT and *mad2*. 2D movies were recorded with a frame rate of 20 seconds. The playback speed is 10 frames per second.

**Movie EV7 Microtubule depolymerization with high concentration of oryzalin causes a SAC-dependent elongation of metaphase I, but no metaphase arrest.** Videos of the representative male meiocytes shown in Fig. 3C. Note how the nucleus is found at the edge of the cell. BMF3 is fused to GFP (green) and TUA5 to TagRFP (magenta). The genotypes and treatments we imaged are from left to right: WT in DMSO and *mad2* in DMSO, WT in 1 µM oryzalin and *mad2* in 1 µM oryzalin. 2D movies were recorded with a frame rate of 2 minutes. The playback speed is 9 frames per second.

**Movie EV8 Meiocytes undergo two NEB events despite the essentially completely depolymerized tubulin.** Videos of the representative male meiocytes shown in Fig. 3C and S3A. Note how the nucleus is found at the edge of the cell before both NEB events. After the second NE reformation, the nucleus “wanders” around in the cell. BMF3 is fused to GFP (green) and TUA5 to TagRFP (magenta). The genotypes and treatments we imaged are from left to right: WT in 1 µM oryzalin and *mad2* in 1 µM oryzalin. 2D movies were recorded with a frame rate of 2 minutes. The playback speed is 10 frames per second.

**Movie EV9 Metaphase I duration in *mps1*, *mad1* and *mad2* is significantly shorter than in the WT.** Videos of representative male meiocytes quantified in Fig. 4A. TUA5 is fused to TagRFP (magenta). The genotypes we analyzed are from left to right: WT, *mps1*, *mad1* and *mad2*. 2D movies were recorded with a frame rate of 30 seconds. The playback speed is 7 frames per second.

**Movie EV10 Metaphase I duration in *bub3.3* is significantly longer than in the WT, contrary to the expected SAC mutant phenotype**. Videos of representative male meiocytes quantified in Fig. 4B. TUA5 is fused to TagRFP (magenta). The genotypes we analyzed are from left to right: WT, *bmf2*, *mad2* and *bub3.3*. 2D movies were recorded with a frame rate of 30 seconds. The playback speed is 10 frames per second.

**Movie EV11 Multiple *bub3.3* cells exhibited severe chromosome alignment defects in metaphase I.** Videos of the representative male meiocytes shown in Fig. 4E. BMF3 is fused to GFP (green) and TUA5 to TagRFP (magenta). The examples shown are from left to right: WT and three *bub3.3* cells with either 0, 1 or multiple mislocalizing chromosome pairs (MC(s)). 2D movies were recorded with a frame rate of 30 seconds. The playback speed is 10 frames per second.

**Movie EV12 SAC double mutants do not proceed faster than SAC single mutants through metaphase I.** Videos of representative male meiocytes quantified in Fig. S4D. TUA5 is fused to TagRFP (magenta). The genotypes we analyzed are from left to right: *bmf2*, *mad1*, *mad2*, *bmf2* x *mad1*, *bmf2* x *mad2* and *mad1* x *mad2*. 2D movies were recorded with a frame rate of 30 seconds. The playback speed is 6 frames per second.

**Movie EV13 The chromosome alignment defect in *bub3.3* is also confirmed with the BMF1 reporter in metaphase I.** Videos of the representative male meiocytes shown in Fig. S4E. BMF1 is fused to GFP (green) and TUA5 to TagRFP (magenta). The examples shown are from left to right: WT and four *bub3.3* cells with either 0, 1, 2 MC at the same pole (s.p.) and 2 MC at different poles (d.p.). 2D movies were recorded with a frame rate of 30 seconds. The playback speed is 10 frames per second.

**Movie EV14 A second T-DNA insertion line, *bub3.3-2*, and the double heterozygous F1 cross *bub3.3* x *bub3.3-2* phenocopy *bub3.3*.** Videos of representative male meiocytes quantified in Fig. S5F-K. BMF3 is fused to GFP (green) and TUA5 to TagRFP (magenta). The genotypes we analyzed are from left to right: WT, *bub3.*3, *bub3.3-2* and *bub3.3* +/− x *bub3.3-2* +/−. 3D movies were recorded with a frame rate of 100 seconds. The playback speed is 5 frames per second.

**Movie EV15 The metaphase I chromosome alignment defect in *bub3.3* is rescued fully by the introgression of a *BUB3.3::gBUB3.3* construct**. Videos of representative male meiocytes quantified in Fig. S5A-E. BMF3 is fused to GFP (green) and TUA5 to TagRFP (magenta). The genotypes we analyzed are from left to right: WT, *bub3.3* and *gBUB3.3* #1 in *bub3.3*. 3D movies were recorded with a frame rate of 100 seconds. The playback speed is 3 frames per second.

**Movie EV16 3D movies of *bub3.3* male meiocytes reveal the full extent of chromosome alignment defects in metaphase I not found in two other SAC mutants, *bmf2* and *mad2***. Videos of the representative male meiocytes shown in Fig. 5A. BMF3 is fused to GFP (green) and TUA5 to TagRFP (magenta). The examples shown are from left to right: WT, *bmf2*, *mad2* and six *bub3.3* cells with either 0 MC, 1 MC, >1 MC at the same spindle pole, >1 MC at different spindle poles, the formation of a perpendicular spindle or the non-rescue of the MC that stays at the spindle pole beyond AO. 3D movies were recorded with a frame rate of 100 seconds. The playback speed is 5 frames per second.

**Movie EV17 Similar chromosome alignment defects are observed in *bub3.3* meiocytes during metaphase II.** Videos of representative male meiocytes quantified in Fig. 6A-F. BMF3 is fused to GFP (green) and TUA5 to TagRFP (magenta). The genotypes we analyzed are from left to right: WT, *bmf2*, *mad2* and *bub3.3*. 3D movies were recorded with a frame rate of 100 seconds. The playback speed is 4 frames per second.

**Movie EV18 *bub3.3* mitotic cells also exhibit severe problems during chromosome alignment.** Videos of representative root epidermal cells shown in Fig. 6G. BMF3 is fused to GFP (green) and TUA5 to TagRFP (magenta). The genotypes we analyzed are from left to right: WT and *bub3.3*. 3D movies were recorded with a frame rate of 1 minute. The playback speed is 5 frames per second.

## References

Alfieri, C., Chang, L., Zhang, Z., Yang, J., Maslen, S., Skehel, M., & Barford, D. (2016). Molecular basis of APC/C regulation by the spindle assembly checkpoint. Nature, 536(7617), 431–436. https://doi.org/10.1038/nature19083

Balboni, M., Yang, C., Komaki, S., Brun, J., & Schnittger, A. (2020). COMET Functions as a PCH2 Cofactor in Regulating the HORMA Domain Protein ASY1. Current Biology, 30(21), 4113–4127.e6. https://doi.org/10.1016/j.cub.2020.07.089

Bomblies, K., Jones, G., Franklin, C., Zickler, D., & Kleckner, N. (2016). The challenge of evolving stable polyploidy: could an increase in “crossover interference distance” play a central role? Chromosoma, 125(2), 287–300. https://doi.org/10.1007/s00412-015-0571-4

Brito, D. A., & Rieder, C. L. (2006). Mitotic Checkpoint Slippage in Humans Occurs via Cyclin B Destruction in the Presence of an Active Checkpoint. Current Biology, 16(12), 1194–1200. https://doi.org/10.1016/j.cub.2006.04.043

Cairo, G., Greiwe, C., Jung, G. I., Blengini, C., Schindler, K., & Lacefield, S. (2023). Distinct Aurora B pools at the inner centromere and kinetochore have different contributions to meiotic and mitotic chromosome segregation. Molecular Biology of the Cell, 34(5), ar43. https://doi.org/10.1091/mbc.E23-01-0014

Cairo, G., MacKenzie, A. M., & Lacefield, S. (2020). Differential requirement for Bub1 and Bub3 in regulation of meiotic versus mitotic chromosome segregation. Journal of Cell Biology, 219(4). https://doi.org/10.1083/jcb.201909136

Capitao, C., Tanasa, S., Fulnecek, J., Raxwal, V. K., Akimcheva, S., Bulankova, P., Mikulkova, P., Khaitova, L. C., Kalidass, M., Lermontova, I., Scheid, O. M., & Riha, K. (2021). A CENH3 mutation promotes meiotic exit and restores fertility in SMG7-deficient Arabidopsis. PLoS Genetics, 17(9), 1–26. https://doi.org/10.1371/journal.pgen.1009779

Comai, L. (2005). The advantages and disadvantages of being polyploid. Nature Reviews Genetics, 6(11), 836–846. https://doi.org/10.1038/nrg1711

Corno, A., Cordeiro, M. H., Allan, L. A., Wei, Q., Harrington, E., Smith, R. J., & Saurin, A. T. (2022). A bifunctional kinase-phosphatase module integrates mitotic checkpoint and error-correction signalling to ensure mitotic fidelity. BioRxiv. https://doi.org/10.1101/2022.05.22.492960

De Jaeger-Braet, J., Krause, L., Buchholz, A., & Schnittger, A. (2021). Heat stress reveals a specialized variant of the pachytene checkpoint in meiosis of Arabidopsis thaliana. The Plant Cell, 2(2), 91–98. https://doi.org/10.1093/plcell/koab257

Demidov, D., Hesse, S., Tewes, A., Rutten, T., Fuchs, J., Karimi Ashtiyani, R., Lein, S., Fischer, A., Reuter, G., & Houben, A. (2009). Aurora1 phosphorylation activity on histone H3 and its cross-talk with other post-translational histone modifications in Arabidopsis. Plant Journal, 59(2), 221–230. https://doi.org/10.1111/j.1365-313X.2009.03861.x

Drpic, D., Almeida, A. C., Aguiar, P., Renda, F., Damas, J., Lewin, H. A., Larkin, D. M., Khodjakov, A., & Maiato, H. (2018). Chromosome Segregation Is Biased by Kinetochore Size. Current Biology, 28(9), 1344–1356.e5. https://doi.org/10.1016/j.cub.2018.03.023

Espert, A., Uluocak, P., Bastos, R. N., Mangat, D., Graab, P., & Gruneberg, U. (2014). PP2A-B56 opposes Mps1 phosphorylation of Knl1 and thereby promotes spindle assembly checkpoint silencing. Journal of Cell Biology, 206(7), 833–842. https://doi.org/10.1083/jcb.201406109

Gorbsky, G. J. (2015). The spindle checkpoint and chromosome segregation in meiosis. FEBS Journal, 282(13), 2458–2474. https://doi.org/10.1111/febs.13166

Hauf, S., Cole, R. W., LaTerra, S., Zimmer, C., Schnapp, G., Walter, R., Heckel, A., Van Meel, J., Rieder, C. L., & Peters, J. M. (2003). The small molecule Hesperadin reveals a role for Aurora B in correcting kinetochore-microtubule attachment and in maintaining the spindle assembly checkpoint. Journal of Cell Biology, 161(2), 281–294. https://doi.org/10.1083/jcb.200208092

Hiruma, Y., Sacristan, C., Pachis, S. T., Adamopoulos, A., Kuijt, T., Ubbink, M., von Castelmur, E., Perrakis, A., & Kops, G. J. P. L. (2015). CELL DIVISION CYCLE. Competition between MPS1 and microtubules at kinetochores regulates spindle checkpoint signaling. Science (New York, N.Y.), 348(6240), 1264–1267. https://doi.org/10.1126/science.aaa4055

Hollister, J. D. (2015). Polyploidy: adaptation to the genomic environment. New Phytologist, 205(3), 1034–1039. https://doi.org/10.1111/nph.12939

Howell, B. J., Moree, B., Farrar, E. M., Stewart, S., Fang, G., & Salmon, E. (2004). Spindle Checkpoint Protein Dynamics at Kinetochores in Living Cells. Current Biology, 14(11), 953–964. https://doi.org/10.1016/j.cub.2004.05.053

Hoyt, M. A., Totis, L., & Roberts, B. T. (1991). S. cerevisiae genes required for cell cycle arrest in response to loss of microtubule function. Cell, 66(3), 507–517. https://doi.org/10.1016/0092-8674(81)90014-3

Izawa, D., & Pines, J. (2015). The mitotic checkpoint complex binds a second CDC20 to inhibit active APC/C. Nature, 517(7536), 631–634. https://doi.org/10.1038/nature13911

Jelluma, N., Brenkman, A. B., van den Broek, N. J. F., Cruijsen, C. W. A., van Osch, M. H. J., Lens, S. M. A., Medema, R. H., & Kops, G. J. P. L. (2008). Mps1 Phosphorylates Borealin to Control Aurora B Activity and Chromosome Alignment. Cell, 132(2), 233–246. https://doi.org/10.1016/j.cell.2007.11.046

Ji, Z., Gao, H., Jia, L., Li, B., & Yu, H. (2017). A sequential multi-target Mps1 phosphorylation cascade promotes spindle checkpoint signaling. ELife, 6, 1–23. https://doi.org/10.7554/eLife.22513

Ji, Z., Gao, H., & Yu, H. (2015). Kinetochore attachment sensed by competitive Mps1 and microtubule binding to Ndc80C. Science, 348(6240), 1260–1264. https://doi.org/10.1126/science.aaa4029

Komaki, S., & Schnittger, A. (2017). The Spindle Assembly Checkpoint in Arabidopsis Is Rapidly Shut Off during Severe Stress. Developmental Cell, 43(2), 172–185.e5. https://doi.org/10.1016/j.devcel.2017.09.017

Komaki, S., Takeuchi, H., Hamamura, Y., Heese, M., Hashimoto, T., & Schnittger, A. (2020). Functional Analysis of the Plant Chromosomal Passenger Complex. Plant Physiology, 183(4), 1586–1599. https://doi.org/10.1104/pp.20.00344

Kozgunova, E., Nishina, M., & Goshima, G. (2019). Kinetochore protein depletion underlies cytokinesis failure and somatic polyploidization in the moss Physcomitrella patens. ELife, 8, 1–16. https://doi.org/10.7554/eLife.43652

Krüger, F. (2017). Vacuole biogenesis in Arabidopsis thaliana. [Ruperto-Carola University of Heidelberg]. https://doi.org/10.11588/heidok.00023502

Kurihara, D., Matsunaga, S., Kawabe, A., Fujimoto, S., Noda, M., Uchiyama, S., & Fukui, K. (2006). Aurora kinase is required for chromosome segregation in tobacco BY-2 cells. Plant Journal, 48(4), 572–580. https://doi.org/10.1111/j.1365-313X.2006.02893.x

Lara-Gonzalez, P., Pines, J., & Desai, A. (2021). Spindle assembly checkpoint activation and silencing at kinetochores. Seminars in Cell and Developmental Biology, 117, 86–98. https://doi.org/10.1016/j.semcdb.2021.06.009

Lara-Gonzalez, P., Scott, M. I. F., Diez, M., Sen, O., & Taylor, S. S. (2011). BubR1 blocks substrate recruitment to the APC/C in a KEN-box-dependent manner. Journal of Cell Science, 124(24), 4332–4345. https://doi.org/10.1242/jcs.094763

Larsen, N. A., Al-Bassam, J., Wei, R. R., & Harrison, S. C. (2007). Structural analysis of Bub3 interactions in the mitotic spindle checkpoint. Proceedings of the National Academy of Sciences, 104(4), 1201–1206. https://doi.org/10.1073/pnas.0610358104

Li, R., & Murray, A. W. (1991). Feedback control of mitosis in budding yeast. Cell, 66(3), 519–531. https://doi.org/10.1016/0092-8674(81)90015-5

Logarinho, E., Resende, T., Torres, C., & Bousbaa, H. (2008). The human spindle assembly checkpoint protein Bub3 is required for the establishment of efficient kinetochore-microtubule attachments. Molecular Biology of the Cell, 19(4), 1798–1813. https://doi.org/10.1091/mbc.e07-07-0633

London, N., & Biggins, S. (2014). Signalling dynamics in the spindle checkpoint response. Nature Reviews Molecular Cell Biology, 15(11), 735–747. https://doi.org/10.1038/nrm3888

London, N., Ceto, S., Ranish, J. A., & Biggins, S. (2012). Phosphoregulation of Spc105 by Mps1 and PP1 regulates Bub1 localization to kinetochores. Current Biology, 22(10), 900–906. https://doi.org/10.1016/j.cub.2012.03.052

Magidson, V., He, J., Ault, J. G., O’Connell, C. B., Yang, N., Tikhonenko, I., McEwen, B. F., Sui, H., & Khodjakov, A. (2016). Unattached kinetochores rather than intrakinetochore tension arrest mitosis in taxol-treated cells. Journal of Cell Biology, 212(3), 307–319. https://doi.org/10.1083/jcb.201412139

Malureanu, L. A., Jeganathan, K. B., Hamada, M., Wasilewski, L., Davenport, J., & van Deursen, J. M. (2009). BubR1 N Terminus Acts as a Soluble Inhibitor of Cyclin B Degradation by APC/CCdc20 in Interphase. Developmental Cell, 16(1), 118–131. https://doi.org/10.1016/j.devcel.2008.11.004

Meraldi, P., Draviam, V. M., & Sorger, P. K. (2004). Timing and checkpoints in the regulation of mitotic progression. Developmental Cell, 7(1), 45–60. https://doi.org/10.1016/j.devcel.2004.06.006

Molè-Bajer, J. (1958). Cine-micrographic analysis of c-mitosis in endosperm. Chromosoma, 9(1), 332–358. https://doi.org/10.1007/BF02568085

Nebel, B. R., & Ruttle, M. L. (1938). The cytological and genetical significance of colchicine. Journal of Heredity, 29(1), 3–10. https://doi.org/10.1093/oxfordjournals.jhered.a104406

Overlack, K., Primorac, I., Vleugel, M., Krenn, V., Maffini, S., Hoffmann, I., Kops, G. J. P. L., & Musacchio, A. (2015). A molecular basis for the differential roles of Bub1 and BubR1 in the spindle assembly checkpoint. ELife, 4(4), 1–24. https://doi.org/10.7554/eLife.05269

Panchy, N., Lehti-Shiu, M., & Shiu, S. H. (2016). Evolution of gene duplication in plants. Plant Physiology, 171(4), 2294–2316. https://doi.org/10.1104/pp.16.00523

Peterson, R., Slovin, J. P., & Chen, C. (2010). A simplified method for differential staining of aborted and non-aborted pollen grains. International Journal of Plant Biology, 1(2), 66–69. https://doi.org/10.4081/pb.2010.e13

Primorac, I., Weir, J. R., Chiroli, E., Gross, F., Hoffmann, I., van Gerwen, S., Ciliberto, A., & Musacchio, A. (2013). Bub3 reads phosphorylated MELT repeats to promote spindle assembly checkpoint signaling. ELife, 2013(2), 1–20. https://doi.org/10.7554/eLife.01030

Prusicki, M. A., Keizer, E. M., van Rosmalen, R. P., Komaki, S., Seifert, F., Müller, K., Wijnker, E., Fleck, C., & Schnittger, A. (2019). Live cell imaging of meiosis in Arabidopsis thaliana. ELife, 8, 1–31. https://doi.org/10.7554/elife.42834

Rieder, C. L., & Maiato, H. (2004). Stuck in Division or Passing through. Developmental Cell, 7(5), 637–651. https://doi.org/10.1016/j.devcel.2004.09.002

Roscioli, E., Germanova, T. E., Smith, C. A., Embacher, P. A., Erent, M., Thompson, A. I., Burroughs, N. J., & McAinsh, A. D. (2020). Ensemble-Level Organization of Human Kinetochores and Evidence for Distinct Tension and Attachment Sensors. Cell Reports, 31(4), 107535. https://doi.org/10.1016/j.celrep.2020.107535

Roy, B., Han, S. J. Y., Fontan, A. N., Jema, S., & Joglekar, A. P. (2022). Aurora B phosphorylates Bub1 to promote spindle assembly checkpoint signaling. Current Biology: CB, 32(1), 237–247.e6. https://doi.org/10.1016/j.cub.2021.10.049

Santaguida, S., Vernieri, C., Villa, F., Ciliberto, A., & Musacchio, A. (2011). Evidence that Aurora B is implicated in spindle checkpoint signalling independently of error correction. EMBO Journal, 30(8), 1508–1519. https://doi.org/10.1038/emboj.2011.70

Saurin, A. T., Van Der Waal, M. S., Medema, R. H., Lens, S. M. A., & Kops, G. J. P. L. (2011). Aurora B potentiates Mps1 activation to ensure rapid checkpoint establishment at the onset of mitosis. Nature Communications, 2(1). https://doi.org/10.1038/ncomms1319

Schindelin, J., Arganda-Carreras, I., Frise, E., Kaynig, V., Longair, M., Pietzsch, T., Preibisch, S., Rueden, C., Saalfeld, S., Schmid, B., Tinevez, J. Y., White, D. J., Hartenstein, V., Eliceiri, K., Tomancak, P., & Cardona, A. (2012). Fiji: An open-source platform for biological-image analysis. Nature Methods, 9(7), 676–682. https://doi.org/10.1038/nmeth.2019

Shah, J. V, Botvinick, E., Bonday, Z., Furnari, F., Berns, M., & Cleveland, D. W. (2004). Dynamics of Centromere and Kinetochore Proteins. Current Biology, 14(11), 942–952. https://doi.org/10.1016/j.cub.2004.05.046

Shepperd, L. A., Meadows, J. C., Sochaj, A. M., Lancaster, T. C., Zou, J., Buttrick, G. J., Rappsilber, J., Hardwick, K. G., & Millar, J. B. A. A. (2012). Phosphodependent recruitment of Bub1 and Bub3 to Spc7/KNL1 by Mph1 kinase maintains the spindle checkpoint. Current Biology, 22(10), 891–899. https://doi.org/10.1016/j.cub.2012.03.051

Sofroni, K., Takatsuka, H., Yang, C., Dissmeyer, N., Komaki, S., Hamamura, Y., Böttger, L., Umeda, M., & Schnittger, A. (2020). CDKD-dependent activation of CDKA;1 controls microtubule dynamics and cytokinesis during meiosis. The Journal of Cell Biology, 219(8). https://doi.org/10.1083/jcb.201907016

Soltis, P. S., & Soltis, D. E. (2009). The role of hybridization in plant speciation. Annual Review of Plant Biology, 60, 561–588. https://doi.org/10.1146/annurev.arplant.043008.092039

Stucke, V. M., Silljé, H. H. W., Arnaud, L., & Nigg, E. A. (2002). Human Mps1 kinase is required for the spindle assembly checkpoint but not for centrosome duplication. EMBO Journal, 21(7), 1723–1732. https://doi.org/10.1093/emboj/21.7.1723

Sudakin, V., Chan, G. K. T., & Yen, T. J. (2001). Checkpoint inhibition of the APC/C in HeLa cells is mediated by a complex of BUBR1, BUB3, CDC20, and MAD2. Journal of Cell Biology, 154(5), 925–936. https://doi.org/10.1083/jcb.200102093

Taylor, S. S., Ha, E., & McKeon, F. (1998). The Human Homologue of Bub3 Is Required for Kinetochore Localization of Bub1 and a Mad3/Bub1-related Protein Kinase. Journal of Cell Biology, 142(1), 1–11. https://doi.org/10.1083/jcb.142.1.1

Tulin, F., & Cross, F. R. (2014). A microbial avenue to cell cycle control in the plant superkingdom. Plant Cell, 26(10), 4019–4038. https://doi.org/10.1105/tpc.114.129312

Van Der Waal, M. S., Saurin, A. T., Vromans, M. J. M., Vleugel, M., Wurzenberger, C., Gerlich, D. W., Medema, R. H., Kops, G. J. P. L., & Lens, S. M. A. (2012). Mps1 promotes rapid centromere accumulation of Aurora B. EMBO Reports, 13(9), 847–854. https://doi.org/10.1038/embor.2012.93

Vanoosthuyse, V., Meadows, J. C., van der Sar, S. J. A., Millar, J. B. A., & Hardwick, K. G. (2009). Bub3p Facilitates Spindle Checkpoint Silencing in Fission Yeast. Molecular Biology of the Cell, 20(24), 5096–5105. https://doi.org/10.1091/mbc.e09-09-0762

Vleugel, M., Omerzu, M., Groenewold, V., Hadders, M. A., Lens, S. M. A., & Kops, G. J. P. L. (2015). Sequential Multisite Phospho-Regulation of KNL1-BUB3 Interfaces at Mitotic Kinetochores. Molecular Cell, 57(5), 824–835. https://doi.org/10.1016/j.molcel.2014.12.036

Vleugel, M., Tromer, E., Omerzu, M., Groenewold, V., Nijenhuis, W., Snel, B., & Kops, G. J. P. L. (2013). Arrayed BUB recruitment modules in the kinetochore scaffold KNL1 promote accurate chromosome segregation. Journal of Cell Biology, 203(6), 943–955. https://doi.org/10.1083/jcb.201307016

Wang, X., Babu, J. R., Harden, J. M., Jablonski, S. A., Gazi, M. H., Lingle, W. L., de Groen, P. C., Yen, T. J., & van Deursen, J. M. A. (2001). The Mitotic Checkpoint Protein hBUB3 and the mRNA Export Factor hRAE1 Interact with GLE2p-binding Sequence (GLEBS)-containing Proteins. Journal of Biological Chemistry, 276(28), 26559–26567. https://doi.org/10.1074/jbc.M101083200

Yang, Y., Tsuchiya, D., & Lacefield, S. (2015). Bub3 promotes Cdc20-dependent activation of the APC/C in S. cerevisiae. Journal of Cell Biology, 209(4), 519–527. https://doi.org/10.1083/jcb.201412036

Younis, A., Hwang, Y.-J., & Lim, K.-B. (2014). Exploitation of induced 2n-gametes for plant breeding. Plant Cell Reports, 33(2), 215–223. https://doi.org/10.1007/s00299-013-1534-y

Zhang, G., Kruse, T., López-Méndez, B., Sylvestersen, K. B., Garvanska, D. H., Schopper, S., Nielsen, M. L., & Nilsson, J. (2017). Bub1 positions Mad1 close to KNL1 MELT repeats to promote checkpoint signalling. Nature Communications, 8(June). https://doi.org/10.1038/ncomms15822

Zhang, H., Deng, X., Sun, B., Lee Van, S., Kang, Z., Lin, H., Lee, Y. R. J., & Liu, B. (2018). Role of the BUB3 protein in phragmoplast microtubule reorganization during cytokinesis. Nature Plants, 4(7), 485–494. https://doi.org/10.1038/s41477-018-0192-z

